# Type I toxin-antitoxin systems contribute to mobile genetic elements maintenance in *Clostridioides difficile* and can be used as a counter-selectable marker for chromosomal manipulation

**DOI:** 10.1101/2020.03.04.976019

**Authors:** Johann Peltier, Audrey Hamiot, Julian R. Garneau, Pierre Boudry, Anna Maikova, Louis-Charles Fortier, Bruno Dupuy, Olga Soutourina

**Affiliations:** Laboratoire Pathogenèse des Bactéries Anaérobies, Institut Pasteur, 75724 Paris Cedex 15, France; Université Paris Diderot, Sorbonne Paris Cité, 75724 Paris Cedex 15, France; Université de Sherbrooke, Faculty of medicine and health sciences, Department of microbiology and infectious diseases, 3201 rue Jean Mignault, Sherbrooke, QC, J1E 4K8, Canada; Center for Data-Intensive Biomedicine and Biotechnology, Skolkovo Institute of Science and Technology, Moscow, 143028, Russia; Université Paris-Saclay, CEA, CNRS, Institute for Integrative Biology of the Cell (I2BC), 91198, Gif-sur-Yvette, France; Institut Universitaire de France (IUF)

**Keywords:** toxin-antitoxin, small noncoding RNA, *cis*-antisense RNA, prophage stability

## Abstract

Toxin-antitoxin (TA) systems are widespread on mobile genetic elements as well as in bacterial chromosomes. According to the nature of the antitoxin and its mode of action for toxin inhibition, TA systems are subdivided into different types. The first type I TA modules were recently identified in the human enteropathogen *Clostridioides* (formerly *Clostridium*) *difficile*. In type I TA, synthesis of the toxin protein is prevented by the transcription of an antitoxin RNA during normal growth. Here, we report the characterization of five additional type I TA systems present within phiCD630-1 and phiCD630-2 prophage regions of *C. difficile* 630. Toxin genes encode 34 to 47 amino acid peptides and their ectopic expression in *C. difficile* induces growth arrest. Growth is restored when the antitoxin RNAs, transcribed from the opposite strand, are co-expressed together with the toxin genes. In addition, we show that type I TA modules located within the phiCD630-1 prophage contribute to its stability and mediate phiCD630-1 heritability. Type I TA systems were found to be widespread in genomes of *C. difficile* phages, further suggesting their functional importance. We have made use of a toxin gene from one of type I TA modules of *C. difficile* as a counter-selectable marker to generate an efficient mutagenesis tool for this bacterium. This tool enabled us to delete all identified toxin genes within the phiCD630-1 prophage, thus allowing investigation of the role of TA in prophage maintenance. Furthermore, we were able to delete the large 49 kb phiCD630-2 prophage region using this improved procedure.

## INTRODUCTION

*Clostridioides difficile* is a medically important human enteropathogen that became a key public health concern over the last two decades in industrialized countries ^1, 2^. This strictly anaerobic spore-forming Gram-positive bacterium is a major cause of antibiotic-associated nosocomial diarrhoea in adults ^3^. The main virulence factors of *C. difficile* are two toxins, TcdA and TcdB, produced by all toxigenic strains ^4^. A binary toxin CDT is also present in some isolates, as well as additional factors that help colonization, like adhesins, pili, and flagella ^5^. Many questions remain unanswered regarding the success of this pathogen and its adaptation within the phage-rich gut environment.

*C. difficile* genome sequencing revealed the mosaic nature of its chromosome, which is composed of more than 10 % of mobile genetic elements including integrated bacteriophages (prophages). Recent studies revealed a high prevalence of prophages in *C. difficile* genomes, each genome harbouring between one and up to five prophages, either integrated into the chromosome or maintained as stable extrachromosomal circular DNA elements ^6^. For example, the largely used laboratory strain 630 carries two homologous prophages, phiCD630-1 and phiCD630-2, while the epidemic ribotype 027 strain R20291 carries one prophage (phi-027). The importance of prophages in the evolution and virulence of many pathogenic bacteria has been clearly demonstrated ^7^. In *C. difficile*, all phages identified so far are temperate and can adopt a lysogenic lifecycle, and some of them have been shown to contribute to virulence-associated phenotypes. This includes modulation of toxin production and complex crosstalk between bacterial host and phage regulatory circuits ^6, 7, 8^. When integrated into *C. difficile* genomes, prophages are stably maintained and replicated along with the host chromosome. However, when they are excised, either spontaneously or following induction by antibiotics or the exposure to other stress conditions, prophages can sometimes be lost during cell division and segregation. The rate of spontaneous phage loss under natural conditions has been estimated to range between 10^−5^ for phage P1 to < 10^−6^ for phage lambda ^9, 10^.

TA modules are widespread in bacteria and archaea. These loci comprise two genes encoding a stable toxin and an unstable antitoxin ^11^. Overexpression of the toxin has either bactericidal or bacteriostatic effects on the host cell while the antitoxin is able to neutralize the toxin action. For all identified TA modules, the toxin is always a protein. The RNA or protein nature and the mode of action of the antitoxin led to the classification of TA modules into six types ^11^. In type I systems, the antitoxin is a small antisense RNA targeting toxin mRNA for degradation and/or inhibition of translation, while in type III systems, the antitoxin RNA binds directly to the toxin protein for neutralization ^12, 13^. For other TA types, both the toxin and the antitoxin are proteins. In most studied type II TA systems, the proteinaceous antitoxin forms a complex with its cognate toxin leading to toxin inactivation ^14^. Major functions of TA modules include plasmid maintenance, abortive phage infection and persister cell formation ^15, 16, 17, 18, 19, 20^. TA loci are commonly found on mobile genetic elements, in particular plasmids in which they were initially discovered and extensively studied. However, the roles of chromosomally-encoded TA modules, including those within prophage genomes, remain largely unexplored. Likewise, the individual contribution from each of the chromosomal TA modules in bacteria remains to be investigated.

We recently reported the identification of the first type I TA systems associated with CRISPR arrays in *C. difficile* genomes ^21^. The co-localization and co-regulation by the general stress response Sigma B factor and biofilm-related factors suggested a possible genomic link between these cell dormancy and adaptive immunity systems. Interestingly, two of these functional type I TA pairs are located within the homologous phiCD630-1 and phiCD630-2 prophages in *C. difficile* strain 630. In the present work, we describe the identification of additional type I TA modules highly conserved within *C. difficile* prophages and provide experimental evidence of their contribution to prophage maintenance and stability.

The unique properties of type I TA systems offer interesting possibilities for biotechnological and therapeutic applications. We demonstrate in the present work that inducible toxicity caused by type I toxins largely improve the efficiency of allele exchange genome editing procedures by promoting the elimination of plasmid-bearing cells.

Altogether, our data provide important insights into the function and possible applications of type I TA systems in *C. difficile*.

## RESULTS

### Identification of novel type I TA pairs in *C. difficile*

Multiple TA modules have been discovered in bacterial chromosomes including prophage regions ^11^. In *C. difficile*, we have recently identified several type I TA pairs adjacent to CRISPR arrays, two of them being located inside the phiCD630-1 and phiCD630-2 prophages of the strain 630 (*CD0956.2*-RCd10 and *CD2907.1*-RCd9, respectively). To determine whether other type I TA modules might be present within phiCD630-1, we performed a bioinformatics analysis on the phiCD630-1 sequence. Due to the small size of the toxin-encoding genes, standard methods of open reading frame (ORF) detection and gene annotation can hinder the identification of all toxin homologs. Moreover, prophages are characterized by a very high gene density, which can impede such detection of small and often overlapping coding regions. We therefore used the tBlastn program (protein query to search against translated genomes) using a previously identified type I toxin CD0956.2 as a query, as it can overcome annotation and ORF detection defects. We identified gene *CD0977.1* and two other novel putative genes, unannotated on the genome, that we named *CD0904.1* and *CD0956.3*. These genes code for small proteins of 47, 35 and 34 amino acids, respectively (Fig. 1A). Prophages phiCD630-1 and phiCD630-2 share a large region of homology with almost identical sequences, which include a duplication of *CD0977.1* and *CD0956.3* (Fig. 1B). In contrast, *CD0904.1* is unique to phiCD630-1 and no other toxin gene homolog could be identified within phiCD630-2. Transcript reads were detected in regions of these putative genes by RNA-seq ^22^ (Fig. S1A and S1B) and a consensus RBS sequence (AGGAGG) was present 7-8 nucleotides upstream of the respective ATG start codons, suggesting that the corresponding proteins might be produced. In addition, all three putative proteins carried a hydrophobic *N*-terminal region and a positively charged tail, which are characteristic features of type I toxins (Fig. 1A). Analysis of our previous TSS mapping data ^22^ and sequence alignments (Fig. S1C) suggested the presence of potential antisense RNAs of these toxin-encoding genes with the presence of TSS associated with Sigma A- and Sigma B-dependent promoter elements in both sense and antisense directions (Fig. S1). Antitoxins of *CD0977.1, CD0904.1* and *CD0956.3*, located on phiCD630-1, were hereafter named RCd11, RCd13 and RCd14, respectively, and those of *CD2889* and *CD2907.2*, found in phiCD630-2, were named RCd12 and RCd15.

**Figure 1.**
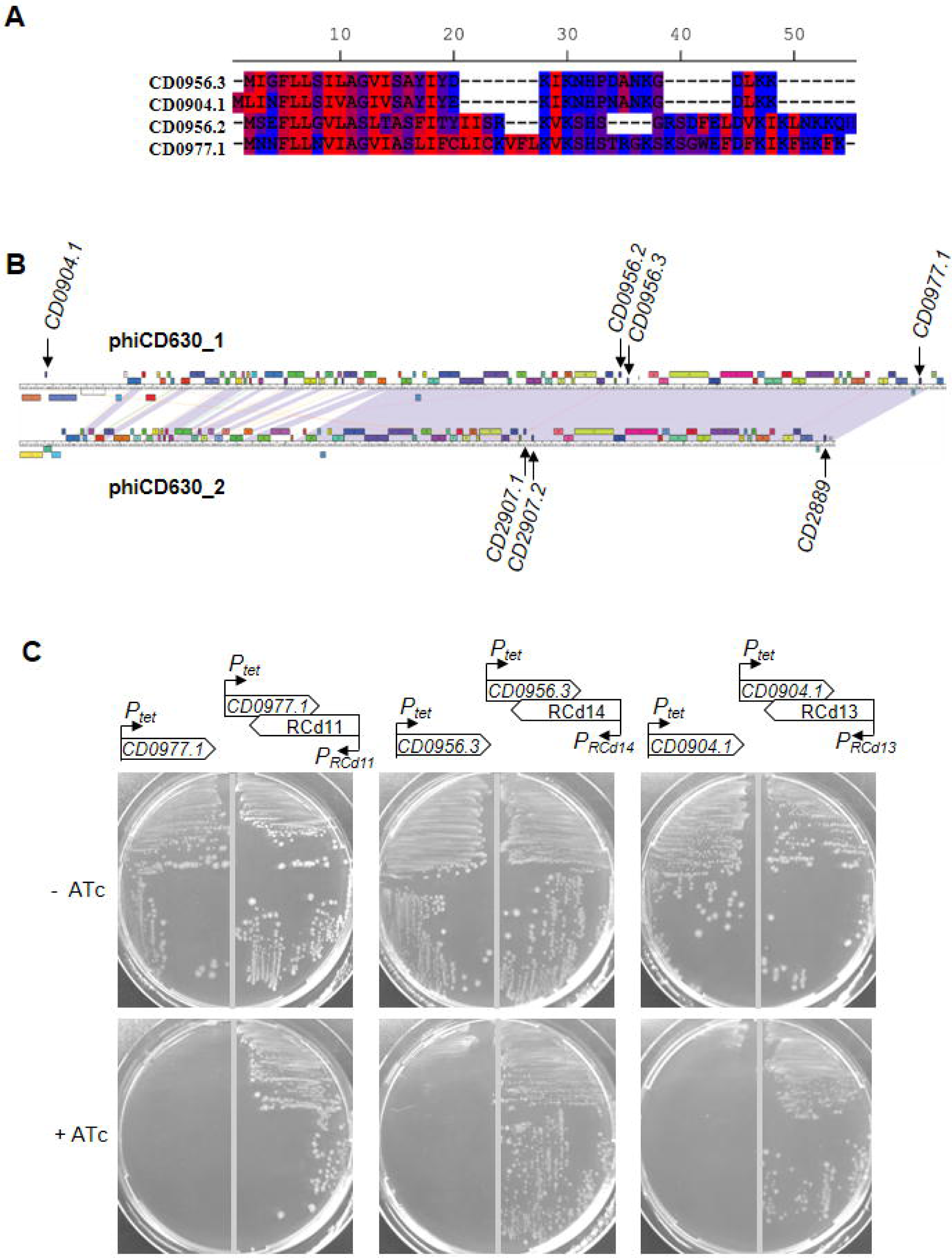
Identification and functionality of novel toxin genes within phiCD630-1. (A) Protein alignment of toxins CD0977.1 with the newly identified CD0904.1, CD0956.2 and CD0956.3. The hydrophobic amino acids are indicated in red. (B) Maps and alignment of the phiCD630-1 and phiCD630-2 genomes. The location of toxin genes in both prophages is indicated. (C) Growth of *C. difficile* 630Δ*erm* strains harbouring the pRPF185-based plasmids on BHI agar plates supplemented with Tm and with (+ATc) or without 10 ng/ml of ATc inducer (- ATc) after 24 hrs of incubation at 37°C. Schematic representations of the constructs are shown.

To determine whether these novel potential TA pairs are functional, pRPF185-derivatives with anhydrotetracycline (ATc)-inducible *P*_*tet*_ promoter were constructed to overexpress *CD0904.1, CD0956.3* and *CD0977.1* toxin genes (pT) or toxin-antitoxin modules (pTA) in *C. difficile* 630Δ*erm*. Antisense RNAs are expressed from their own promoter in pTA. Growth of 630Δ*erm* carrying the different pT and pTA vectors on BHI plates was indistinguishable in the absence of ATc inducer (Fig. 1C). In contrast, growth of the 630Δ*erm*/pT strains was completely inhibited when ATc was present in the medium, while strains 630Δ*erm/*pTA showed a reversion of the growth defect. These results demonstrate that *CD0904.1, CD0956.3* and *CD0977.1* encode potent toxins and are associated with antisense RNAs that function as antitoxins.

### Detailed characterization of the *CD0977.1*-RCd11 TA pair

Intriguingly, predicted riboswitches responding to the c-di-GMP signalling molecule, cdi1_4 and cdi1_5, precede RCd11 and RCd12 antisense RNAs ^22^. We therefore sought to further characterize the *CD0977.1*-RCd11 TA pair. In agreement with the data above, addition of ATc to liquid cultures in exponential growth phase led to an immediate growth arrest of strain 630Δ*erm*/pT, unlike 630Δ*erm*/p (Fig. S2A). In addition, the growth arrest was accompanied by a drop of colony-forming units (CFUs) (Fig. S2B). Similarly to previous observations with other *C. difficile* type I TA modules ^21^, the analysis of liquid cultures by light microscopy showed that toxin overexpression was accompanied by an increase in cell length in about 10% of the cells (Fig. S2C). Their length was above the mean length value of 630Δ*erm*/p control strain with two standard deviations (10.5 µm). The co-expression of the entire TA module led to the partial reversion of this phenotype.

Using Northern blotting, we detected both toxin and antitoxin transcripts in the 630Δ*erm*/p, 630Δ*erm*/pT (*CD0977.1*) and 630Δ*erm*/pTA (*CD0977.1*-RCd11) strains (Fig. 2A). In the absence of ATc inducer, a major transcript of about 300 nt was detected in all three strains with a *CD0977.1-*specific probe. When using a RCd11-specific probe, transcripts of about 150, 300 and 400 nt were observed. Under inducing conditions, a reverse correlation between the relative toxin and antitoxin transcript abundance was noticed. To determine the impact of c-di-GMP on the antitoxin transcripts, we elevated c-di-GMP intracellular levels in the 630Δ*erm* wild type strain by expressing the diguanylate cyclase gene *dccA* from a plasmid (p*dccA*) (Fig. 2B), as previously reported ^22^. A c-di-GMP-regulated read-through transcript of about 400 nt, as well as a terminated transcript of about 140 nt were detected in this strain by Northern blotting using a riboswitch-specific probe. In contrast, abundance of toxin transcripts was not affected by fluctuations of c-di-GMP levels.

**Figure 2.**
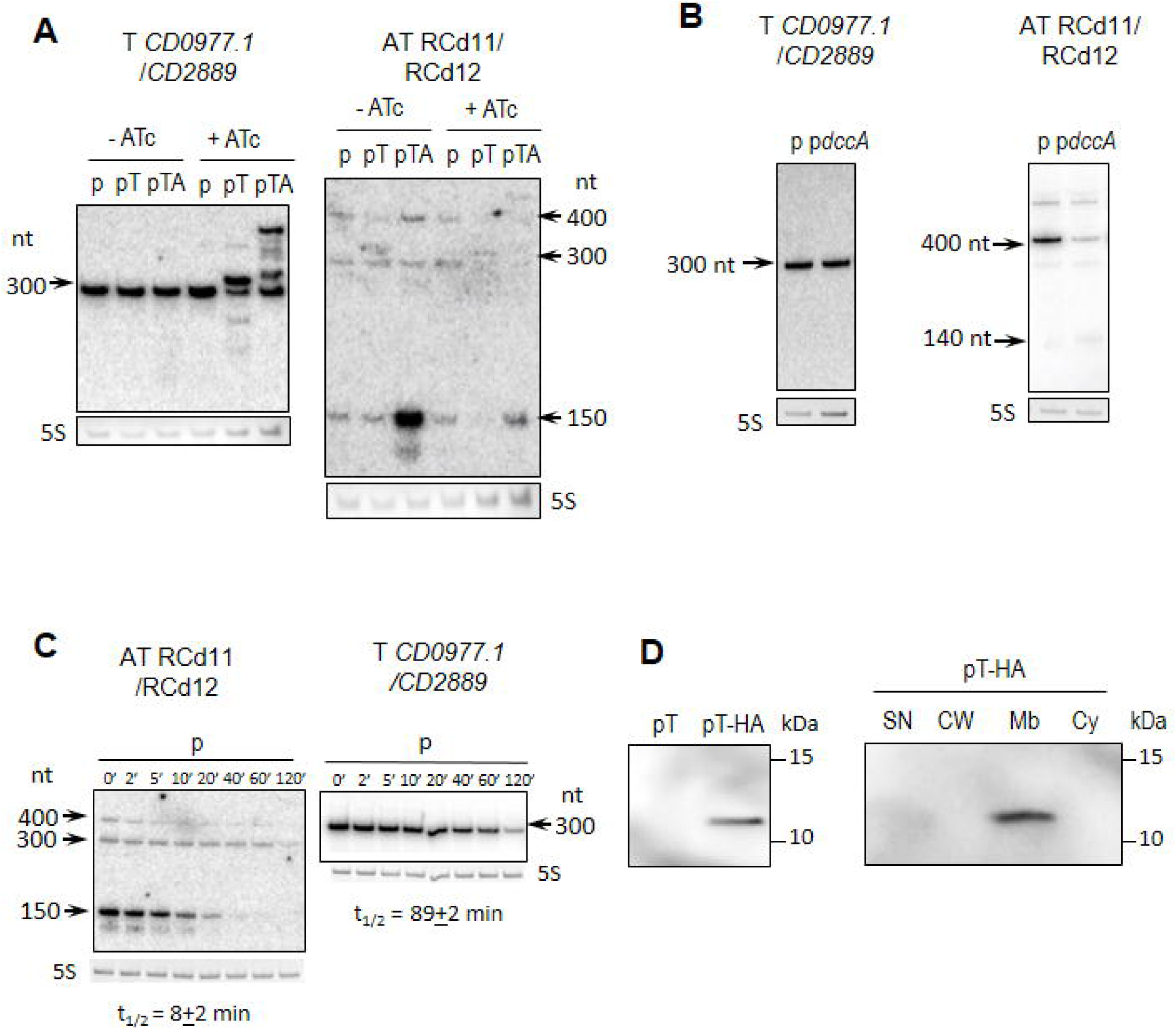
Detection of *CD0977.1* and RCd11 transcripts and CD0977.1-HA protein. (A) Northern blot of total RNA from *C. difficile* carrying p (empty vector), pT (expression of *CD0977.1*) or pTA (expression of *CD0977.1* and its antitoxin) in absence (- ATc) or in presence (+ ATc) of 250 ng/ml of the inducer ATc. (B) Northern blot of total RNA from *C. difficile* carrying p (empty vector) or p*dccA* (expression of the diguanylate cyclase encoding gene *dccA*) in presence of 250 ng/ml ATc. (C) Northern blot of total RNA from *C. difficile* 630Δ*erm* carrying an empty vector (wt/p) collected at the indicated time after addition of rifampicin. All Northern blots were probed with a radiolabelled oligonucleotide specific to the toxin (T CD0977.1/CD2889) or the antitoxin (AT RCd11/RCd12) transcript and 5S RNA at the bottom serves as loading control. The arrows show the detected transcripts with their estimated size. The relative intensity of the bands was quantified using the ImageJ software. (D) Detection and subcellular localization of the CD0977.1-HA protein. Immunoblotting with anti-HA detected a major polypeptide of ∼12 kDa in whole cell extracts of *C. difficile* carrying pT-HA (CD0977.1-HA) grown in presence of 250 ng/ml of ATc but not in extracts of *C. difficile* carrying pT (non-tagged CD0977.1) (left panel). The culture of *C. difficile* carrying pT-HA was fractionated into supernatant (SN), cell wall (CW), membrane (Mb) and cytosolic (Cy) compartments and immunoblotted with anti-HA antibodies. Proteins were separated on 12% Bis-Tris polyacrylamide gels in MES buffer.

We then mapped the transcriptional start (TSS) and termination sites for the genes of the potential RCd11/RCd12-*CD0977.1/CD2889* TA modules by 5’/3’RACE analysis (Fig. S3, Table S3). The results obtained agreed generally well with the transcript lengths deduced from TSS mapping, RNA-seq and Northern blot. Taken together, these data suggest the presence of two tandem TSS for RCd11, i.e. *P*_*1*_ associated with c-di-GMP-dependent riboswitch and *P*_*2*_ located downstream from the riboswitch (Fig. S1).

It is in the nature of type I antitoxins to be short-lived in contrast to the stable toxin mRNA ^13^. To determine the half-lives of toxin and antitoxin RNAs of the *CD0977.1*-RCd11 module, *C. difficile* strains were grown in TY medium until late-exponential phase and rifampicin was added to block transcription. Samples were taken at different time points after rifampicin addition for total RNA extraction and Northern blot analysis with toxin and antitoxin-specific probes. In a control strain 630Δ*erm*/p carrying an empty vector, the half-life of major 150-nt antitoxin transcript for RCd11 was estimated to be about 8 min while the half-life of *CD0977.1* toxin mRNA was estimated to about 89 min (Fig. 2C). Interestingly, depletion of the RNA chaperone protein Hfq resulted in a moderate destabilization of *CD0977.1* toxin mRNA and antitoxin RCd11 RNA with the half-life of 64 min and 4 min, respectively (Fig. S4). By contrast, the stable *CD0977.1* toxin mRNA was further stabilized to over 120 min half-life in the strains depleted for RNase III, RNase J and RNase Y. For antitoxin RCd11 RNA, we also observed a stabilization in strain depleted for RNase Y (Fig. S4) suggesting that this ribonuclease could contribute to antitoxin RNA degradation.

To confirm the protein nature of CD0977.1 and assess its subcellular localization, we constructed a derivative of CD0977.1 with an HA tag fused to the C-terminus of CD0977.1 expressed from a plasmid under the control of the inducible *P*_*tet*_ promoter (pT-HA). *C. difficile* strain carrying pT-HA was grown to mid-exponential phase, induced with ATc for 90 min, and whole cell extracts were prepared. Induction of *CD0977.1* expression immediately stopped the growth, as revealed by OD_600_ measurements (data not shown), suggesting that the HA-tag does not interfere with toxin activity. HA-tagged CD0977.1 was detectable by Western blotting with anti-HA antibodies (Fig. 2D). No band was observed in a whole cell extract of a control strain producing untagged CD0977.1 protein. The distribution of HA-tagged CD0977.1 within supernatant, cell wall, membrane and cytosolic compartments was studied (Fig. 2D). HA-tagged CD0977.1 was only detected in the membrane fraction, indicating the association of CD0977.1 with the cell membrane of *C. difficile*.

### The antitoxin transcript controlled by cdi1_5 riboswitch is dispensable for efficient toxin inactivation

To get further insights into the function of abundant short (transcribed from *P*_*2*_) and less abundant long RCd11 antitoxin transcripts (transcribed from *P*_*1*_), we generated new plasmid constructs that allowed the inducible expression of the *CD0977.1* toxin gene under the control of the *P*_*tet*_ promoter and the expression of different forms of the RCd11 antisense RNA (Fig. 3A). The first construct, yielding pDIA6816, lacked the cdi1_5 riboswitch and its associated promoter (*P*_*1*_) but retained the *P*_*2*_ promoter of the antitoxin. On the opposite, the second construct, yielding pDIA6817, retained the *P*_*1*_ promoter and the associated riboswitch but had a disrupted *P*_*2*_ promoter. The construct in which both promoters of RCd11 were present (pDIA6785) served as a positive control for the assay, and the RCd11 promoterless construct (pDIA6335) was included as a negative control. All plasmids were introduced into *C. difficile* 630Δ*erm* and the corresponding strains were grown on BHI plates supplemented with 10 and 100 ng/ml ATc to induce *CD0977.1* toxin expression. Growth of the strain carrying pDIA6816 was similar to that observed for the control strain carrying pDIA6785 in the presence of 10 ng/ml ATc and was slightly defective in the presence of 100 ng/ml ATc (Fig. 3A and Fig. S5A). In contrast, the strain carrying pDIA6817 did not grow in the presence of 10 or 100 ng/ml ATc, similarly to the negative control strain. Similar results were obtained when the strains were grown in an automatic plate reader for 20 h in liquid medium in the presence of 5 ng/ml ATc (Fig 3A). Interestingly, induction of toxin expression on BHI plate with a lower dose of ATc (5 ng/ml) led to a partial reversion of the growth defect of the strain carrying pDIA6816, unlike the negative control strain (Fig. S5A). These results suggest that the short antitoxin transcript driven by promoter *P*_*2*_ is crucial for the efficient inactivation of the toxin, while the longer antitoxin transcript directed by *P*_*1*_ is dispensable.

**Figure 3.**
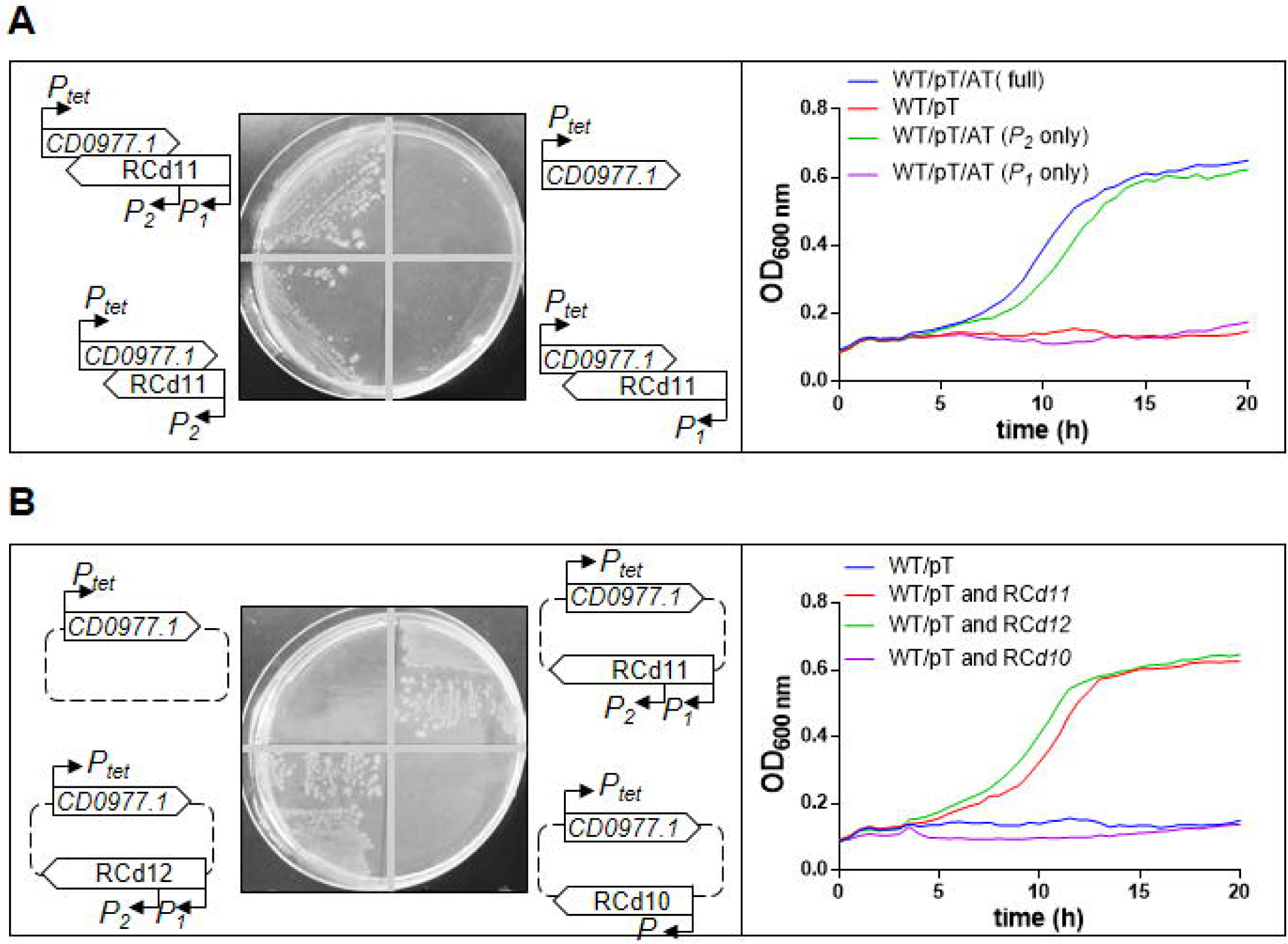
Impact of toxin-antitoxin co-expression on growth. The effect on the toxicity of *CD0977.1* of long and short antitoxin transcripts expressed *in cis* (A) and *in trans* (B) was assessed. Growth of *C. difficile* 630Δ*erm* strains harbouring the pRPF185-based plasmids on BHI agar plates supplemented with Tm and 10 ng/mL of ATc inducer after 24 hrs of incubation at 37°C and in TY broth at 37°C in the presence of 5 ng/mL ATc. Schematic representations of the constructs are shown.

### RCd12 counteracts toxic activity of non-cognate CD0977.1 toxin

Nucleotide sequences of short RCd11, lying within phiCD630-1 and short RCd12, lying within phiCD630-2, are almost identical with only 3 mismatches located near the 3’ end, in the region overlapping with the toxin transcript (Fig. S6A and S7). We therefore wondered whether RCd12 could cross-react with the transcript of the non-cognate toxin CD0977.1. To answer this question, we generated new constructs in which *CD0977.1* toxin gene under the control of the *P*_*tet*_ promoter and different antitoxins with their own promoter were co-expressed from the same plasmid but from distant locations (Fig. 3B). As anticipated, expression of RCd11 in *trans* counteracted the toxicity associated with the expression of the cognate toxin both on plate and in liquid culture (Fig. 3B and Fig. S5B). Replacement of RCd11 with RCd12 led to the same result. By contrast, expression of the more divergent RCd10 (the antitoxin of CD0956.2 toxin from a previously characterized TA module lying within phiCD630-1^21^) in *trans* failed to revert the growth defect induced by *CD0977.1* expression (Fig. S5B and 3B). These data indicate that antitoxins act in a highly specific manner to repress their cognate toxins, not only when they are expressed from the native convergent TA configuration, but also when expressed in *trans*. However, the specificity of interaction is permissive for at least 3 mismatches allowing RCd12 expressed from phiCD630-2 to efficiently prevent CD0977.1 toxin production from phiCD630-1.

### TA modules confer plasmid stabilization

TA systems have been initially discovered on plasmids where they confer maintenance of the genetic element ^17^. Plasmid loss results in a rapid decrease in the levels of the unstable antitoxin, which allows the stable toxin to inhibit cell growth. To test whether the TA modules located on phiCD630-1 could contribute to plasmid maintenance, we assessed the stability of plasmids in which the toxin genes and their cognate antitoxin were co-expressed under the control of their respective native promoter in *C. difficile* 630Δ*erm*. Bacteria harbouring the empty vector pMTL84121 were used as a control. After 7 passages in TY broth in the absence of antibiotic pressure, pMTL84121 was maintained by 1.0% (+/-0.4%) of the bacterial population (Fig. 4). In contrast, plasmids expressing TA pairs were still present in 22.3 (+/- 5.4%) to 63.6 % (+/- 14.3%) of total cells. These results indicate that the four TA pairs can confer plasmid maintenance.

**Figure 4.**
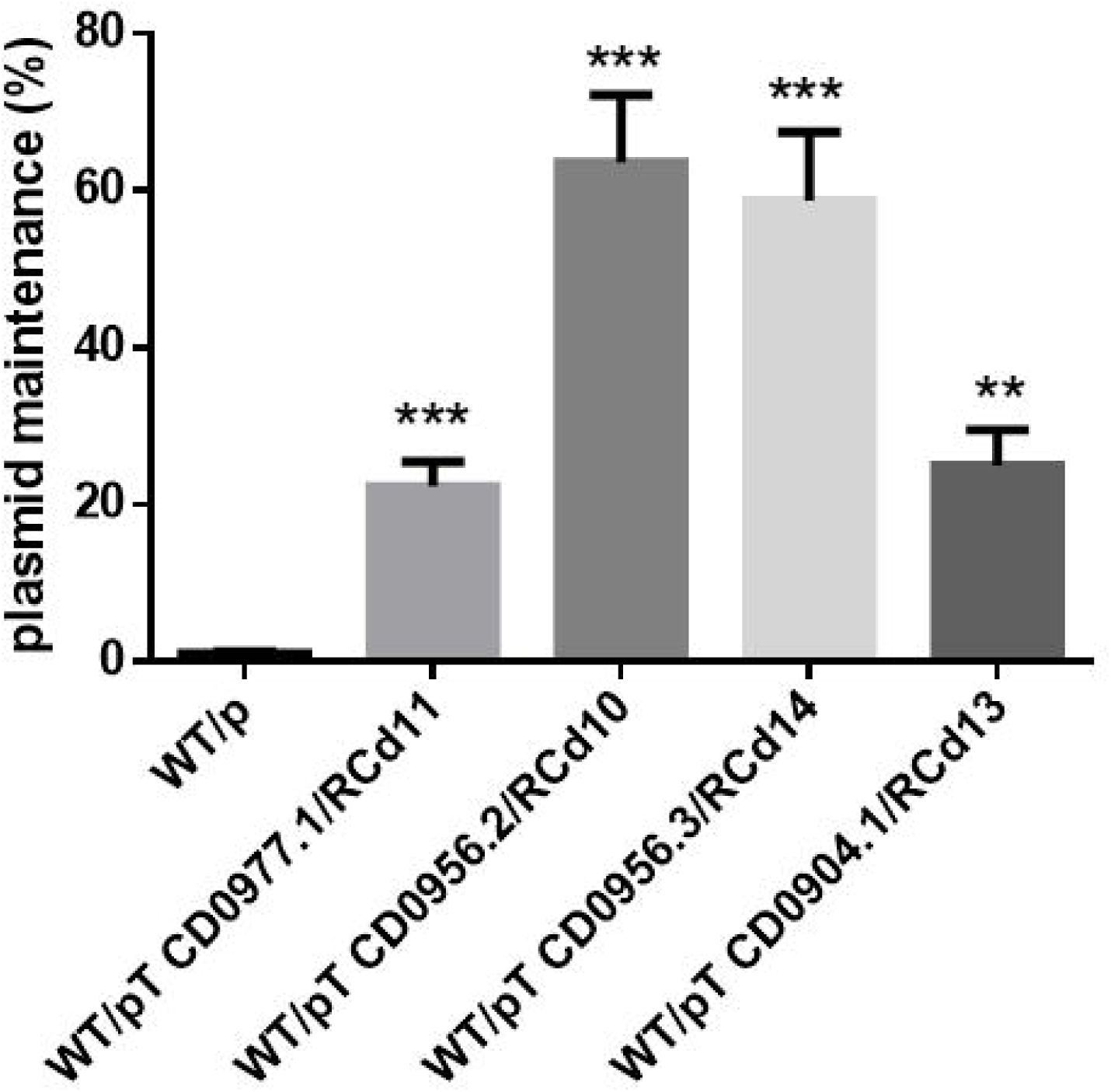
Impact of toxin-antitoxin modules on plasmid loss in the absence of selection pressure. The stability of pMTL84121 (p, empty vector) and pMTL84121-derived vectors expressing the different TA modules in *C. difficile* 630Δ*erm* was determined after seven passages (every 12 hours) in TY broth without thiamphenicol. Values represent means ± standard deviations (*N* = 3). ** *P* ≤ 0.05 and *** *P* ≤ 0.001 by an unpaired *t* test.

### Deletion of phiCD630-1 toxin genes in *C. difficile* 630Δerm

In order to determine whether the TA modules contribute to phiCD630-1 stability, we undertook the construction of deletion mutants of toxin genes in *C. difficile* 630Δ*erm*. For this purpose, we constructed a new Allele-Coupled Exchange (ACE) vector, derived from pMTL-SC7315, a *codA*-based “pseudosuicide” plasmid that replicates at a rate lower than that of the host chromosome ^23^. The *codA* cassette was here replaced with the *CD2517.1* toxin gene placed under the control of the *P*_*tet*_ inducible promoter (Fig. S8A). We demonstrated the functionality of RCd8-*CD2517.1* type I TA module in *C. difficile* in a previous study ^21^. In our new vector, designated pMSR, the inducible toxic expression of *CD2517.1* is used as a counter-selection marker to screen for plasmid excision and loss (see Materials and Methods) and this greatly facilitates the isolation of *C. difficile* deletion mutant generated by double cross-over allele exchange (Fig. S8C and D). We also constructed a second vector, pMSR0, for allele exchange in *C. difficile* ribotype 027 strains and other ribotypes (see Materials and Methods and Fig S8B). By using this new tool, we first deleted the 49.3 kb phiCD630-2 locus to prevent any interfering cross-talk with phiCD630-1 (Fig. S9). A multiple deletion mutant of toxin-encoding genes *CD0904.1, CD0956.2, CD0956.3* and *CD0977.1* (ΔT4) was then generated in the ΔphiCD630-2 background (Table S1).

### TA systems are involved in maintenance of phiCD630-1 in the host cells

Because the loss of an integrated phage from cells first requires its excision from the host genome, we sought to determine whether spontaneous excision of phiCD630-1 from chromosomal DNA occurred. To do so, we performed a PCR on genomic DNA from *C. difficile* 630Δ*erm* ΔphiCD630-2 with primers flanking the *attL* and *attR* sites of phiCD630-1 (Fig. S10A). A PCR product with a size of 88 bp corresponding to the excised prophage was detected (Fig. S10B) and DNA sequencing of this amplicon confirmed the complete removal of phiCD630-1 from the host chromosome. A second PCR-based assay showed that the excised prophage (PCR product of 117 bp) was present as an extrachromosomal circular form in the host cell (Fig. S10A and B). We could also deduce the *attB, attP, attL* and *attR* sites from the sequencing of the PCR products (Fig. S10C). However, the frequency of phiCD630-1 excision, measured by quantitative PCR (qPCR), was very low (∼ 0.015 %) (Fig. S11). To screen for the presence/absence of phiCD630-1 in the host cells, we introduced, using our new ACE vector, the *ermB* gene placed under the control of the strong *thl* promoter of *Clostridium acetobutylicum* as previously described ^24^, into an innocuous location (between *CD0946.1* and *CD0947*) of phiCD630-1 and phiCD630-1-ΔT4 (Fig. 5A). Starting from the overnight cultures, the ΔphiCD630-2 phiCD630-1::*erm* and ΔphiCD630-2 phiCD630-1-ΔT4::*erm* strains were subcultured four times in fresh medium and cells were screened for erythromycin resistance by plating onto non-selective and erythromycin-containing agar plates. Nearly 100% of cells from both strains were found to still be resistant to erythromycin in these conditions, indicating that they had retained the prophage (Fig S10D).

**Figure 5.**
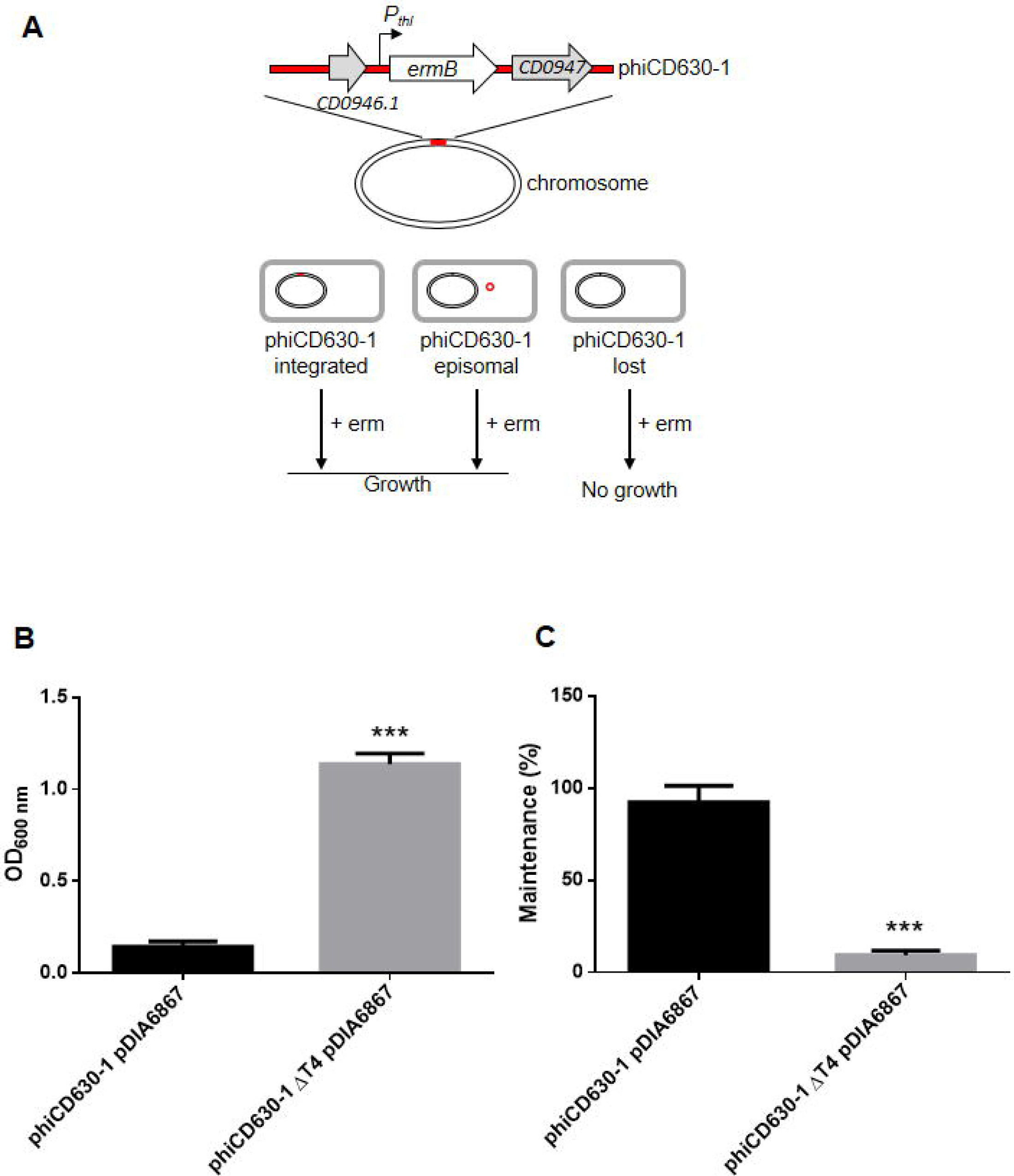
Impact of toxin /antitoxin modules on prophage maintenance. (A) Schematic representation of the method used to quantify prophage maintenance. A cassette containing an erythromycin resistance gene (*ermB*) under control of the strong *thl* promoter of *C. acetobutylicum* was introduced into an innocuous location of phiCD630-1, within the intergenic region between *CD0946.1* and *CD0947* genes, encoding a hypothetical protein and a putative scaffold protein, respectively. Cultures grown for 24 hrs were plated on erythromycin-containing agar plates and cells that lost the prophage were selectively killed. (B) and (C): Strains ΔphiCD630-2 phiCD630-1::*erm* and ΔphiCD630-2 phiCD630-1-ΔT4::*erm* carrying a pDIA6867 (overproducing the excisionase CD0912) were inoculated at an initial optical density at 600 nm (OD_600nm_) of 0.005 in TY medium supplemented with 7.5 μg/ml Tm and 10 ng/ml ATc. Cultures were incubated at 37°C for 24 hrs and bacterial growth was determined by measurement of the OD_600nm_ (B) and the maintenance of prophages was quantified by plating serial dilutions on agar plates supplemented or not with 2.5 μg/ml Erm (C). Values represent means ± standard deviations (*N* = 3). *** *P* ≤ 0.001 by an unpaired *t* test.

In an attempt to artificially increase the excision rate of phiCD630-1, we ectopically expressed the putative excisionase *CD0912* from the inducible *P*_*tet*_ promoter, yielding pDIA6867. CD0912, identified in a bioinformatics search, is a 109 amino acid protein with a predicted DNA-binding domain similar to the HTH-17 superfamily and the excisionase (Xis) family ^25^. Induction of *CD0912* expression with 10 ng/ml ATc in *C. difficile* 630Δ*erm* resulted in a high excision rate of phiCD630-1 (∼ 70 %), indicating that CD0912 functions as an excisionase for phiCD630-1 (Fig. S11). Expression of *CD0912* in strains ΔphiCD630-2 phiCD630-1::*erm* and ΔphiCD630-2 phiCD630-1ΔT4::*erm* caused excision at a similar rate, suggesting that TA systems do not affect phiCD630-1 excision (Fig. S11).

Strains ΔphiCD630-2 phiCD630-1::*erm* and ΔphiCD630-2 phiCD630-1-ΔT4::*erm* carrying pDIA6867 were then inoculated at an initial OD_600_ of 0.005 in TY supplemented with 7.5 μg/ml Tm and 10 ng/ml ATc. After 24 hrs of incubation at 37°C, measurement of the OD_600_ revealed a dramatic growth defect of ΔphiCD630-2 phiCD630-1::*erm* compared to the ΔphiCD630-2 phiCD630-1-ΔT4::*erm* (Fig. 5B). In addition, plating of the cells bearing pDIA6867 on non-selective and erythromycin-containing agar plates revealed that phiCD630-1::*erm* was still present in more than 90% of the total population while phiCD630-1-ΔT4::*erm* remained in less than 10 % of the cells (Fig. 5C). These results thus show that TA modules are important for phiCD630-1 maintenance after its excision and highlight the impact of the toxin expression on the cell growth upon the loss of prophage. Together, these data demonstrate that TA modules mediate phiCD630-1 heritability.

### Type I TA are prevalent in *C. difficile* phage genomes

Since we identified additional toxin variants of type I TA systems after careful inspection of phiCD630-1 full genome, we decided to re-scan for possible ORFs in every available phage genomes of *C. difficile* using the permissive algorithm of the NCBI ORFfinder software. ORFs with minimal length of 60 nucleotides as well as nested ORFs were detected. A blastP search against the corresponding proteins allowed the identification of toxin homologs in all *C. difficile* prophage genomes (functional phages) (Fig. 6). Moreover, toxin sequence alignments revealed the high conservation of the hydrophobic *N*-terminal region, as well as the lysine-rich, positively charged region at the C-terminus. Hence, these data suggest the functionality of the toxins and reinforce their proposed role for phage maintenance and preservation.

**Figure 6.**
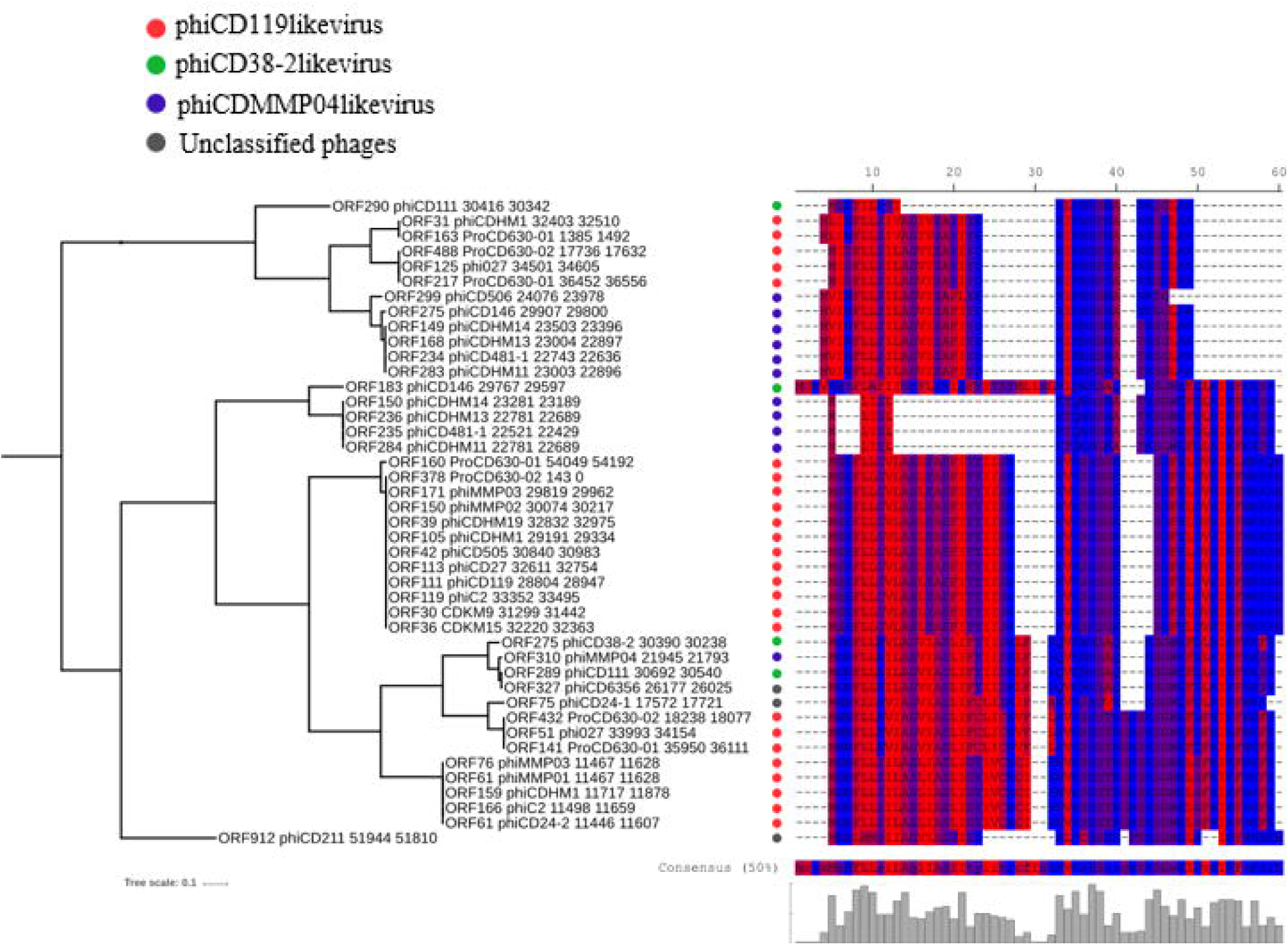
Relationship between phage phylogeny and toxin variants. Putative toxin protein sequences detected in all available phage genomes were aligned using MUSCLE (v3.8) algorithm (EMBL-EBI). All phage genomes were re-scanned for potential ORFs using the NCBI ORFfinder software and detected ORFs were translated into their corresponding protein sequence. Protein sequences were combined to create a local BLASTp database and confirmed functional toxins CD0956.2 (large variant) and CD0904.1 (short variant) were used as queries to search for putative toxins in the database. Significant hits (min 45% identity, min 28% coverage) were retrieved and the corresponding proteins were aligned using MUSCLE (v3.8) algorithm (EMBL-EBI). The protein sequence consensus is shown. A phylogenetic tree was built using the Poisson distance method and neighbour joining implemented in Seaview (v4.4.2). Residues were coloured according to high hydrophobicity (red) and low hydrophobicity (blue).

Despite these conserved regions, alignment of toxins also revealed small variations among sequences. We therefore sought to explore the possible relationship between phage phylogeny and the observed toxin variants. A whole genome comparison of all phages included in this study was performed to create phylogenetic groups (phiCD119-like viruses, phiCD38-2-like viruses and phiMMP04-like viruses), as previously described ^26^. A clear link between phage groups and specific toxin variants could be established, suggesting an independent acquisition of TA systems in different groups of phages (Fig. 6 and Fig. S12). Interestingly, an extended search outside *C. difficile* phages revealed the presence of other toxin homologs inside plasmids of *C. difficile* and *Paeniclostridium sordellii*, a closely related species (Fig. S13). These findings imply that *C. difficile* phages could recombine with plasmids to exchange genetic material, as already proposed for *E. coli* phages ^27, 28^.

## DISCUSSION

In this study, we identified and characterized novel functional type I TA modules in *C. difficile* 630 prophages. Although these modules share characteristic features of TA systems, i.e. (i) toxic proteins are membrane-associated and have a positively charged tail, (ii) the toxin mRNA is much more stable than the antitoxin RNA, (iii) artificial expression of the toxin genes inhibits bacterial growth unless their cognate antitoxin RNA is co-expressed; they do not present sequence homology with other TA modules identified in other bacteria.

RCd11-*CD0977.1* and RCd12-*CD2889* TA pairs are duplications respectively located within the homologous phiCD630-1 and phiCD630-2 prophages. Two alternative RCd11 (and RCd12) antitoxin transcripts were identified and the longer, less abundant form is associated with the cdi1_4 (and cdi1_5) c-di-GMP-responsive riboswitch. C-di-GMP is a second messenger in bacterial systems and a key signal in the control of critical lifestyle choices, such as the transition between planktonic and biofilm growth ^22, 29^. c-di-GMP has been found to regulate important functions in *C. difficile*, including motility, production of type IV pili, cell aggregation and biofilm formation, through control of gene expression by c-di-GMP-dependent riboswitches ^29^. Sixteen predicted c-di-GMP sensing riboswitches are encoded in the *C. difficile* 630 genome and the regulatory function of five of them has been investigated so far. Interestingly, cdi1_4 and cdi1_5 riboswitches were recently reported to be insensitive to an elevation of the c-di-GMP levels, and only transcript reads corresponding to the riboswitch sequences themselves (the short form) were detected, and no read-through seemed to occur ^30^. In contrast, our RACE-PCR and Northern-blot analysis indicated the presence of a transcript downstream from these riboswitches. Moreover, our data suggested that cdi1_4 and cdi1_5 are functional riboswitches responding to c-di-GMP since the abundance of the downstream transcripts was significantly reduced in the presence of high levels of c-di-GMP. Despite these data, we found that the shorter and more abundant RCd11 transcript alone was sufficient to ensure complete CD0977.1 toxin inactivation under our conditions. In contrast, the longer antitoxin transcript associated with the cdi1_4 riboswitch could only counteract the CD0977.1 toxicity when the toxin gene was expressed at low levels. This suggests that this antitoxin transcript might be involved in the tight regulation of CD0977.1 production and might be crucial to prevent toxin translation under conditions where expression levels of the toxin gene would be slightly higher than those of the short antitoxin transcript. The c-di-GMP levels would then be critical in this regulation since elevated levels would result in a decreased abundance of the short RCd11 transcript and consequently in growth inhibition.

In this work, we detected a natural background excision of the phiCD630-1 prophage and we identified the phage excisionase gene, *CD0912*. Expression of *CD0912* from a plasmid promoted high levels of prophage excision from the host chromosome. While phiCD630-1 and phiCD630-2 of *C. difficile* 630 share a large region of duplicated sequence, it is worth noting that *CD0912* is located in the variable region and has no homolog in phiCD630-2. Interestingly, no obvious putative excisionase-encoding gene could be identified in phiCD630-2 although natural excision of this prophage could also be detected in the course of our experiments. Moreover, expression of *CD0912* had no impact on the excision rate of phiCD630-2. This suggests that phiCD630-2 might encode an atypical, yet to be identified excisionase. TA systems have been shown to play three important biological functions, i.e., stabilization of mobile genetic elements (post-segregational killing), abortive phage infection and persister cell formation ^31^. Prophage maintenance is among the suggested function of TA including a recent example of type II TA system that stabilizes prophage in *Shewanella oneidensis* ^32^ and another type II TA system promoting the maintenance of an integrative conjugative element in *Vibrio cholerae* ^33^. This physiological function in prophage stabilization was also suggested for type I TA modules but had never been experimentally demonstrated prior to this study ^34^. Prophage excision upon expression of *CD0912* made phiCD630-1 prone to be lost by the host cells and we could thus show that type I TA systems are important to maintain the episomal form of the phage into the host cell. Indeed, the frequency of phiCD630-1ΔT4 loss was higher than that of wildtype phiCD630-1. Moreover, maintenance of phiCD630-1 in the cells expressing the excisionase gene was associated with a strong growth defect, which can be attributed to the post-segregational killing mechanism. We can speculate that the unstable antitoxin is degraded in daughter cells where the phage has been lost and the stable toxic protein prevents the growth of the new cell. Expression of the excisionase gene in cells carrying the prophage devoid of the toxin genes also resulted in a growth defect, although to a lesser extent. This suggests that a supplementary TA system might be present in phiCD630-1 or that the excisionase has an additional function affecting the cell growth and that remains to be uncovered. Several experimental conditions were tested in this work to induce the loss of the phage from the cells. Surprisingly, four passages of the strain carrying phiCD630-1 with the intact toxin genes in rich medium with constant expression of the excisionase gene resulted in approximately 99% of loss of this prophage. This is likely due to a progressive enrichment of the cell population surviving the loss of the phage since the growth rate of this population is higher than that of the population bearing the phage. In any case, these data suggest that the identification and the overexpression of the phage excisionase-encoding genes could provide an easy and efficient way to cure *C. difficile* strains from their prophages.

Toxins of type I TA systems are relatively small proteins, and this is probably one of the reasons why they have remained uncharacterized and unexplored in *C. difficile* and in other organisms. In this study, we have come to realize that standard methods of annotation are unable to detect all toxin homologs present in prophage genomes. Novel toxin homologs, previously unannotated, could thus be detected inside plasmids of *C. difficile* and *P. sordellii*. It has been proposed that phages could recombine with plasmids during infection of the same or different bacterial species to exchange genetic material ^8, 27, 28^. It is therefore tempting to speculate that this TA system could originate from *P. sordellii*, or at least that it has the ability to disseminate, through horizontal gene transfer involving conjugation and recombination, from one species to another. Intriguingly, it was previously noticed that a 1.9-kb region could have been transferred from the plasmid of a *C. difficile* 630 strain to the phiCD38-2 prophage ^8^. It was suggested that this recombination event had led to the acquisition of *parA*, a gene assumed to help the newly created chimeric phage to autonomously replicate and segregate as a circular plasmid. Our analysis of TA systems in phiCD38-2 shows that this 1.9-kb region also carried a TA (gp33) that presumably contributes to the phage maintenance and stability. It is interesting to observe that TA encoding regions can relocate from one mobile genetic element to another in this fashion, and that genes in proximity to the TA being transferred (i.e. *parA* gene) have more chances to become fixed in the newly integrated DNA. In the latter case, the region transferred seems to provide two complementary and beneficial features to the phage, i.e. the capacity to segregate successfully to the daughter cell, and the death of the cells upon curing if the phage has not been sequestered in both dividing cells. However, since TA systems behave as selfish elements that promote their propagation within bacterial genomes at the expense of their host ^35, 36^, they are likely to be maintained and observed after their transfer by recombination events, even if they bring no selective advantage.

Besides their biological functions, TA modules are also versatile tools for a multitude of purposes in basic research and biotechnology ^37^. For example, the MazF toxin-encoding gene from *E. coli* is used as a counter-selection marker for chromosomal manipulation in *Bacillus subtilis* and *C. acetobutylicum* ^38, 39^. In this study, we engineered an inducible counter-selection marker based on the *C. difficile CD2517.1* toxin gene. When *CD2517.1*, cloned under the control of an inducible promoter on a plasmid, is expressed in *C. difficile* upon addition of the inducer, bacterial growth is immediately interrupted ^21^. Taking advantage of this feature, we generated a novel vector for allele exchange in *C. difficile* 630 and another vector for use in *C. difficile* ribotype 027 strains and other ribotypes strains. It should be noted that expression of the RCd8 antitoxin from the pMSR0 vector was required to counteract the basal expression of *CD2517.1* toxin gene due to the *P*_*tet*_ leakiness. In contrast, expression of the RCd8 antitoxin from the pMSR vector was not required since the *CD2517.1*-RCd8 TA module is naturally present within the chromosome of *C. difficile* 630. Native expression of RCd8 was therefore sufficient to prevent CD2517.1 production from the plasmid. The system developed by Cartman *et al.*, which uses the *codA* gene coding for cytosine deaminase as a counter-selection marker for allelic exchange mutations was appealing at first since no genomic manipulations were needed to apply the counter-selection ^23^. However, *codA*-based counter-selection was somewhat ineffective in our hands and false-positive counter-selected colonies with the plasmid still integrated into the chromosome were repeatedly found. This was reported by the authors as the consequence of loss-of-function mutations in either *codA* or *upp* genes leading to the bypass of the counter-selection. Our newly developed system proved to be much more efficient than all the others we have tested so far, and we did not observe any false-positive clones so far. We successfully used this system to construct multiple mutants in various *C. difficile* strains, including the ΔT4 mutant lacking the four toxin genes reported in this study, deletion of a large chromosomal region of 50 kb corresponding to the phiCD630-2 prophage, and gene insertion into the bacterial chromosome (*ermB* gene). We therefore expect these new vectors to become invaluable genetic tools that will foster research in *C. difficile*.

## MATERIALS AND METHODS

### Plasmids, bacterial strains construction and growth conditions

*C. difficile* and *Escherichia coli* strains and plasmids used in this study are presented in Table S1. *C. difficile* strains were grown anaerobically (5 % H_2_, 5 % CO_2_, and 90 % N_2_) in TY ^40^ or Brain Heart Infusion (BHI, Difco) media in an anaerobic chamber (Jacomex). When necessary, cefoxitin (Cfx; 25 μg/ml), cycloserine (Cs; 250 μg/ml) and thiamphenicol (Tm; 7.5 μg/ml) were added to *C. difficile* cultures. *E. coli* strains were grown in LB broth, and when needed, ampicillin (100 μg/ml) or chloramphenicol (15 μg/ml) was added to the culture medium. The non-antibiotic analogue anhydrotetracycline (ATc) was used for induction of the *P*_*tet*_ promoter of pRPF185 vector derivatives in *C. difficile* ^41^. Strains carrying pRPF185 derivatives were generally grown in TY medium in the presence of 250 ng/ml ATc and 7.5 µg/ml Tm for 7 h, unless stated otherwise. Growth curves were obtained using a GloMax plate reader (Promega).

All primers used in this study are listed in Table S2. For expression of TA genes under the control of their native promoters, the *CD0977.1*-RCd11, *CD0956.2*-RCd10, *CD0904.1*-RCd13, *CD0956.3*-RCd14 TA modules and their associated promoters were amplified by PCR and cloned into BamHI and XhoI sites of pMTL84121 ^42^. For inducible expression of *C. difficile* genes, we used the pDIA6103 derivative of pRPF185 vector expression system lacking the *gusA* gene ^22^. To assess the activity of the putative toxin CD0977.1 and the functionality of its cognate antitoxin, the *CD0977.1* gene (−66 to +222 relative to the translational start site) and the RCd11-*CD0977.1* region (−220 to +608 relative to the translational start site) were amplified by PCR and cloned into StuI and BamHI sites of pDIA6103 vector under the control of the ATc-inducible *P*_*tet*_ promoter; yielding pDIA6335 and pDIA6337. All other constructs described below were performed with NEBuilder HiFi DNA Assembly (NEB). For this purpose, the pDIA6103 vector was linearized by inverse PCR at the MCS for cloning of genes under the control of the ATc-inducible *P*_*tet*_ promoter. To assess the toxicity of CD0904.1 and CD0956.3 and the functionality of their cognate antitoxin, the *CD0904.1* (−43 to + 109 relative to the translational start site) and *CD0956.3* genes (−35 to + 108 relative to the translational start site) as well as the RCd13-*CD0904.1* (−43 to +230 relative to the translational start site) and the RCd14-*CD0956.3* regions (−35 to + 248 relative to the translational start site) were amplified by PCR and cloned into the linearized pDIA6103 vector, yielding pDIA6866, pDIA7030, pDIA6934 and pDIA7031, respectively. For inducible toxin expression and co-expression of its cognate antitoxin, the RCd11-*CD0977.1* TA region with the two RCd11 promoters (−66 to +608 relative to the translational start site of *CD0977.1*) or the RCd11-*CD0977.1* TA region with the *P*_*2*_ promoter only (−66 to +306 relative to the translational start site of *CD0977.1*) were amplified by PCR and cloned into the linearized pDIA6103 vector, yielding pDIA6785 and pDIA6816, respectively. For inducible toxin expression and co-expression of its cognate antitoxin with the promoter *P*_*1*_ only, the identified transcriptional start site (TSS) G of *P*_*2*_ promoter was replaced with a T and the -10 box TATAAT was replaced with TTTTTT, using an inverse PCR approach, to disrupt the *P*_*2*_ promoter; yielding pDIA6817. To investigate the action of cognate (RCd11) and non-cognate (RCd10 and RCd12) antitoxins on toxin CD0977.1 when co-expressed *in trans* (from a site distant from the vector MCS), we used an inverse PCR approach to construct different plasmids on the basis of pDIA6335. RCd11 (−158 to +386 relative to the first TSS), RCd12 (−158 to +386 relative to the TSS) and RCd10 (−167 to +282 relative to the TSS) with their respective promoter were amplified by PCR and cloned into pDIA6335 linearized by inverse PCR at a site distant from the vector MCS, yielding pDIA6791, pDIA6792 and pDIA6793, respectively. For subcellular localization of toxin CD0977.1, we used an inverse PCR approach to construct *CD0977.1*-HA-expressing plasmids (pDIA6622) on the basis of pDIA6335 with primers designed to introduce the hemagglutinin HA-tag sequence at the C-terminal part of toxin coding regions, directly upstream of the stop codon (Table S2). For expression of the CD0912 excisionase, the *CD0912* gene (−37 to + 333 relative to the translational start site) was amplified by PCR and cloned into the linearized pDIA6103 vector, yielding pDIA6867.

The resulting pRPF185 derivative plasmids were transformed into the *E. coli* HB101 (RP4) and subsequently mated with the appropriate *C. difficile* strains (Table S1). *C. difficile* transconjugants were selected by sub-culturing on BHI agar containing Tm (15 µg/ml), Cfx (25 µg/ml) and Cs (250 µg/ml).

### Mutagenesis approach and mutant construction

To improve the efficiency of the allele exchange mutagenesis in *C. difficile*, we made use of the inducible toxicity of the CD2517.1 type I toxin that we previously reported ^21^. To construct the pMSR vector, used for allele exchange in C. difficile 630Δ*erm*, the *codA* gene was removed from the “pseudosuicide” vector pMTL-SC7315 ^23^ by inverse PCR and replaced by a 1169 bp fragment comprising the entire *P*_*tet*_ promoter system and the downstream *CD2517.1* toxin gene. This fragment was amplified from pDIA6319 plasmid ^21^ and the purified PCR product was cloned into the linearized plasmid. In parallel, the pMSR0 vector, for allele exchange in *C. difficile* ribotype 027 strains and other ribotypes, was constructed by removing the *codA* gene from the vector pMTL-SC7215 by inverse PCR and replacing it with the *CD2517.1*-RCd8 TA region with *CD2517.1* under the control of the *P*_*tet*_ promoter, as described above, and RCd8 under the control of its own promoter. For deletions, allele exchange cassettes were designed to have between 800 and 1050 bp of homology to the chromosomal sequence in both up- and downstream locations of the sequence to be altered. The homology arms were amplified by PCR from *C. difficile* strain 630 genomic DNA (Table S1) and purified PCR products were directly cloned into the PmeI site of pMSR vector using NEBuilder HiFi DNA Assembly. To insert *P*_*thl*_-*ermB* into the phiCD630-1 prophage, within the intergenic region between *CD0946.1* and *CD0947* genes, homology arms (∼900 bp up- and downstream of the insertion site) were amplified by PCR from strain 630 genomic DNA (Table S1). The *P*_*thl*_*-ermB* cassette was amplified from the Clostron mutant *cwp19* ^24, 43^. Purified PCR products were all assembled and cloned together into the PmeI site of pMSR vector using NEBuilder HiFi DNA Assembly.

All pMSR-derived plasmids were initially transformed into *E. coli* strain NEB10β and all inserts were verified by sequencing. Plasmids were then transformed into *E. coli* HB101 (RP4) and transferred by conjugation into the appropriate *C. difficile* strains. The adopted protocol for allele exchange was similar to that used for the *codA*-mediated allele exchange ^23^, except that counter-selection was based on the inducible expression of the *CD2517.1* toxin gene. Transconjugants were selected on BHI supplemented with Cs, Cfx and Tm, and then restreaked onto fresh BHI plates containing Tm. After 24 h, faster-growing single-crossover integrants formed visibly larger colonies. One such large colony was restreaked once or twice on BHI Tm plate to ensure purity of the single crossover integrant. Purified colonies were then restreaked onto BHI plates containing 100 ng/ml ATc inducer to select for cells in which the plasmid had been excised and lost. In the presence of ATc, cells in which the plasmid is still present produce CD2517.1 at toxic levels and do not form colonies. Growing colonies were then tested by PCR for the presence of the expected deletion.

### Light microscopy

For light microscopy, bacterial cells were observed at 100x magnification on an Axioskop Zeiss Light Microscope. Cell length was estimated for more than 100 cells for each strain using ImageJ software ^44^.

### Subcellular localization of HA-tagged toxins by cell fractionation and Western blotting

*C. difficile* cultures were inoculated from overnight grown cells in 10 ml of TY medium at an optical density at 600 nm (OD_600_) of 0.05. Cultures were allowed to grow for 3 hours before the addition of 250 ng/ml ATc and incubation for 90 min. Then, cells were centrifuged and proteins were extracted. Cell lysis, fractionation and protein analysis were performed as previously described ^45^. Coomassie staining was performed for loading and fractionation controls. Western blotting was performed with anti-HA antibodies (1:2, 000) (Osenses) using standard methods.

### RNA extraction, Northern blot and 5’/3’RACE

Total RNA was isolated from *C. difficile* strains after 4, 6 or 10 hrs of growth in TY medium, or 7.5 hrs in TY medium containing 7.5 µg/ml of Tm and 250 ng/ml of ATc for strains carrying pRPF185 derivatives, as previously described ^46^. Starvation conditions corresponded to a 1 h incubation of exponentially grown cells (6 h of growth) in PBS buffer at 37°C. Northern blot analysis and 5’/3’RACE experiments were performed as previously described ^22^.

### Measurement of RNA decay by rifampicin assay

For determination of toxin and antitoxin RNA half-lives the *C. difficile* strains were grown in TY medium supplemented with 250 ng/ml ATc and 7.5 µg/ml Tm for 7.5 h at 37°C. Samples were taken at different time points after the addition of 200 µg/mL rifampicin (0, 2, 5, 10, 20, 40, 60 and 120 min) and subjected to RNA extraction and Northern blotting.

### Plasmid stability assays

Overnight cultures of *C. difficile* cells containing the pMTL84121 empty vector or the pMTL84121 derivatives were grown in TY broth with Tm and used to inoculate (at 1%) 5 ml of fresh TY broth without antibiotic. Every 10 to 14 hrs, 1% of the cultures were reinoculated into fresh TY broth without antibiotic. After seven passages, CFUs were estimated on TY plates supplemented or not with Tm to differentiate between the total number of cells and the plasmid-containing cells.

### Quantification of the frequency of prophage excision

The frequency of prophage excision in different *C. difficile* strains was estimated by quantitative PCR on genomic DNA extracted using the NucleoSpin Microbial DNA kit (Macherey-Nagel). The total chromosome copy number was quantified based on the reference gene *dnaF* (*CD1305*) encoding DNA polymerase III. The number of chromosomes devoid of phiCD630-1 was quantified by PCR amplification using primers flanking phiCD630-1 (Table S2), which only results in PCR products when the prophage is excised.

### PhiCD630-1 stability assays

Overnight cultures of *C. difficile* strain 630 ΔphiCD630-2 phiCD630-1::*erm* and ΔphiCD630-2 phiCD630-1-ΔT4::*erm* were used to inoculate 10 ml of TY broth at an initial OD_600_ of 0.05. Every 10 to 14 h, cultures were subcultured at an initial OD_600_ of 0.05. After four passages, the cultures were serially diluted and plated on BHI plates to estimate the total CFUs and on BHI plates supplemented with 2.5 µg/ml erythromycin to determine the number of CFUs in which phiCD630-1 was still present. For cells expressing the excisionase gene, overnight cultures of *C. difficile* strain 630 ΔphiCD630-2 phiCD630-1::*erm* and ΔphiCD630-2 phiCD630-1-ΔT4::*erm* carrying pDIA6867 were used to inoculate fresh TY broth with Tm and 10 ng/ml ATc at an initial OD_600_ of 0.005. After 24 hrs of incubation at 37°C, cultures were serially diluted and plated on BHI plates to estimate the total CFUs and on BHI plates supplemented with 2.5 µg/ml erythromycin to determine the number of CFUs in which phiCD630-1 was still present. The OD_600_ of the cultures was also measured to monitor cell growth.

## ACKNOWLEGEMENTS

This work was supported by Agence Nationale de la Recherche (“CloSTARn”, ANR-13-JSV3-0005-01 to O.S.), the Institut Universitaire de France (to O.S.), the University Paris-Saclay, the Institute for Integrative Biology of the Cell, the Pasteur Institute, the DIM-1HEALTH regional Ile-de-France program (LSP grant no. 164466), the CNRS-RFBR PRC 2019 (grant no. 288426 № 19-54-15003) to O.S., and a Vernadski fellowship to A.M. We would like to thank Marc Monot for helpful discussions.

## AUTHOR CONTRIBUTIONS

J.P. and O.S. conceived and coordinated the study, which was initiated by P.B. J.P. and O.S. performed the majority of the experiments. A.H. constructed vectors and deletion mutants. J.R.G. performed the *in-silico* analyses, A.M. performed growth curves and light microscopy. L-C.F. and B.D. provided scientific insight into the design of the experiments. J.P. and O.S. wrote the paper and all authors reviewed and approved the final version of the manuscript.

## COMPETING INTERESTS

The authors declare that they have no conflict of interest.

## SUPPLEMENTAL FIGURE LEGENDS

**Figure S1. RNA-seq and TSS mapping profiles for the TA loci in *C. difficile* strain 630.** The TAP-/TAP+ profile comparison for 5’-end RNA-seq data is aligned with RNA-seq data for *CD0977.1*-RCd11 (A) and *CD0904.1*-RCd13 TA (B) genomic regions. The TSS are indicated by red broken arrows in accordance with the positions of 5’-transcript ends shown by vertical green lines on the sequence read graphs corresponding to TSS. TSS corresponds to positions with significantly greater numbers of reads in TAP+ samples. 5’-end sequencing data show 51-bp reads matching to the 5’-transcript ends, while RNA-seq data show reads covering the whole transcript. Coding sequences are indicated by blue arrows and the regulatory RNAs are indicated by grey arrows. The promoter regions of the antitoxins are shown. The positions of TSS “+1” are represented. (C) Alignment between the sequences of *CD0904.1*-RCd13 (top line) and *CD0956.3*-RCd14 (bottom line) using EMBOSS Needle. Green boxes show the open reading frames of *CD0904.1* and *CD0956.3* and green arrows indicate the direction of the genes. Blue boxes show the -10 and the -35 boxes of the promoter regions and black boxes show the transcription start site of RCd13 and RCd14. The deduced promoter region of RCd14 antitoxin is represented. In (A), (B) and (C), the red boxes show the Sigma A-dependent promoter -10 and -35 elements and the yellow boxes shows the Sigma B-dependent promoter element.

**Figure S2. Impact of toxin CD0977.1 expression on cell viability and morphology.** Growth (A) and viability (B) of *C. difficile* 630Δ*erm* strain carrying the pRPF185-based plasmids (empty: p or with *CD0977.1* toxin gene under the control of the *P*_*tet*_ promoter: pT) in TY broth in presence of 200 ng/ml ATc. The time point of ATc addition is indicated by a vertical arrow. Values represent means ± standard deviations (*N* = 3). * *P* ≤ 0.05 by a Student’s *t* test. (C) Selected images from light microscopy observations of 630/p, 630/pT and 630/pTA strains grown in TY broth for 1 h at 37°C after the addition of 250 ng/mL ATc. Cell length was estimated using the ImageJ software for at least 115 cells per strain. The mean values with standard deviations are indicated for each strain, as well as the proportion of cells with length above 2 standard deviations relative to the 630/p control strain mean length.

**Figure S3. Sequence of type I *CD0977.1*-RCd11 TA locus in *C. difficile*.** The thick horizontal arrows below the double-stranded sequences show the toxin and antitoxin transcripts and the direction of transcription. The transcriptional start sites for sense and antisense transcripts identified by 5’/3’RACE and TSS mapping are indicated by vertical arrows with their genomic location. Line thickness corresponds to the proportion of observed extremities. The genomic location of 5’- and 3’-ends of the transcripts are indicated above the sequence. The inverted repeats at the position of transcriptional terminators are indicated by thin black arrows. The positions of Sigma A-dependent -10 and -35 promoter elements of antitoxin (AT) are shown in red boxes. The positions of Sigma A-dependent -10 and -35 elements promoter, ribosome binding site, translation initiation codon and stop codon of toxin (T) mRNA are underlined in blue. The positions of Sigma B-dependent promoter elements are shown in green boxes for both TA genes.

**Figure S4. Northern blots showing the stability of *CD0977.1/CD2889* toxin (A) and RCd11/RCd12 antitoxin (B) transcripts in strains depleted for RNase III, RNase Y, RNase J and Hfq.** To determine half-lives, samples were taken at the indicated time points after the addition of 200 µg/mL rifampicin. RNAs were extracted from strains CDIP369 (630/p), CDIP230 expressing an antisense RNA for the *rncS* gene encoding RNase III (AS *rncS*), CDIP53 strain expressing an antisense RNA for the *hfq* gene (AS *hfq*), CDIP55 strain expressing an antisense RNA for the *rnJ* gene encoding RNase J (AS *rnJ*) and CDIP57 strain expressing an antisense RNA for the *rny* gene encoding RNase Y (AS *rny*). All Northern blots were probed with a radiolabelled oligonucleotide specific to the toxin (T CD0977.1/CD2889) or the antitoxin (AT RCd11/RCd12) transcript and 5S RNA at the bottom serves as loading control. The relative intensities of the bands were quantified using ImageJ software.

**Figure S5. Impact of toxin-antitoxin co-expression on growth.** The effect on the toxicity of *CD0977.1* of long and short antitoxin transcripts expressed *in cis* (A) and *in trans* (B) was assessed. Growth of *C. difficile* 630Δ*erm* strains harbouring the pRPF185-based plasmids on BHI agar plates supplemented with Tm and the indicated concentration of ATc inducer after 24 hrs of incubation at 37°C. Schematic representations of the constructs are shown.

**Figure S6. Nucleotide alignment of antitoxins.** (A) Nucleotide alignment of RCd11 and RCd12 using LALIGN. (B) Nucleotide alignment of RCd11 and RCd10 using LALIGN.

**Figure S7. Secondary structure prediction of *CD0977.1* mRNAs and corresponding antitoxin RCd11 RNA.** The RNA secondary structure prediction was performed by Mfold software. Ribosome binding site (SD), translation initiation codon and stop codon positions are highlighted. The positions of mismatches in RCd12 AT sequence are indicated.

**Figure S8. New vector for efficient gene editing in *C. difficile*.** Features of the pMSR (A) and pMSR0 (B) vectors used for allele exchange in *C. difficile* 630Δ*erm* and *C. difficile* 027 ribotype strains, respectively. The toxin gene *CD2517.1* is under the control of the ATc inducible promoter *P*_*tet*_ and the RCd8 antitoxin present in pMSR0 is under control of its own promoter. Schematic overview of the allele exchange protocol (C) and of the inducible counterselection method used to isolate double cross-over clones (D). Isolated single cross-over integrants were restreaked on ATc-containing agar plates to induce synthesis of toxin CD2517.1. Cells that kept the pMSR plasmid (either integrated or excised) produced CD2517.1 and were selectively killed.

**Figure S9. Deletion of the phiCD630-2 prophage from *C. difficile* 630Δ*erm* using the newly developed allele exchange method.** (A) Schematic representation of phiCD630-2 in *C. difficile* 630Δ*erm*. The location of primers used to screen for mutants is represented. (B) PCR products amplified using the indicated primers from the parental strain 630Δ*erm* (WT) and the ΔphiCD630-2 strain (Δ). A product of 1,348 bp could be amplified with primers JP528-JP527 if phiCD630-2 had been deleted, whereas a product of 1,339 bp could be amplified with primers JP570-JP527 if phiCD630-2 was still present.

**Figure S10. Maintenance and site-specific excision of phiCD630-1 from genomic DNA of *C. difficile* 630.** (A) Schematic representation of phiCD630-1 DNA excision from genomic DNA of *C. difficile* 630, and circularization. The location of primers used to demonstrate prophage excision is represented. (B) PCR products amplified using the indicated primers from *C. difficile* 630 genomic DNA. (C) DNA sequences within *attP, attB, attL* and *attR* as determined by Sanger sequencing. The central identical sequences where recombination occurs are shown in bold. Short segments of sequence surrounding the central identity region are shown in blue (bacterial sequences) and in red (phage sequences). (D) The maintenance of prophage in strains ΔphiCD630-2 phiCD630-1::*erm* and ΔphiCD630-2 phiCD630-1-ΔT4::*erm* was determined after four passages in TY broth by plating serial dilutions on agar plates supplemented or not with 2.5 μg/ml Erm. Values represent means ± standard deviations (*N* = 3).

**Figure S11.** Impact of excisionase (CD0912) overproduction on excision of phiCD630-1 and phiCD630-1-ΔT4. Excision rate of phiCD630-1 was higher in *C. difficile* carrying pDIA6867 (inducible expression of *CD0912*) in presence of 10 ng/ml of the inducer ATc when compared to the ΔphiCD630-2 strain without plasmid and was not impacted by the deletion of toxin genes. Values represent means ± standard deviations (*N* = 3).

**Fig. S12. Heatmap and phylogenetic tree showing *C. difficile* relatedness for 27 sequenced *C. difficile* phage genomes available in the NCBI database.** The Gegenees software (v2.2.1) was used to produce the heatmap of genome similarity. Similarity scores are based on a fragmented all-against-all pairwise alignment using BLASTn and the accurate alignment option (fragment size, 200; step size, 100). The colours reflect the similarity, ranging from low (red) to high (green). Phages were assigned to a genus if they clustered closely to another phage previously described as a member of that genus. The phylogenetic tree is based on the sequence similarity scores from the same whole-genome comparison and was constructed using the neighbour joining method with the SplitsTree4 software (v 4.13.1).

**Fig. S13. Identification of toxin homologs outside *C. difficile* phages.** Toxin homologs found outside *C. difficile* phages were aligned. The protein sequence consensus is shown at the bottom. Phylogenic analysis of toxins is also represented.

