## Supplemental figures for "Type I toxin-antitoxin systems contribute to mobile genetic elements maintenance in *Clostridioides difficile* and can be used as a counter-selectable marker for chromosomal manipulation"

### Slide 1
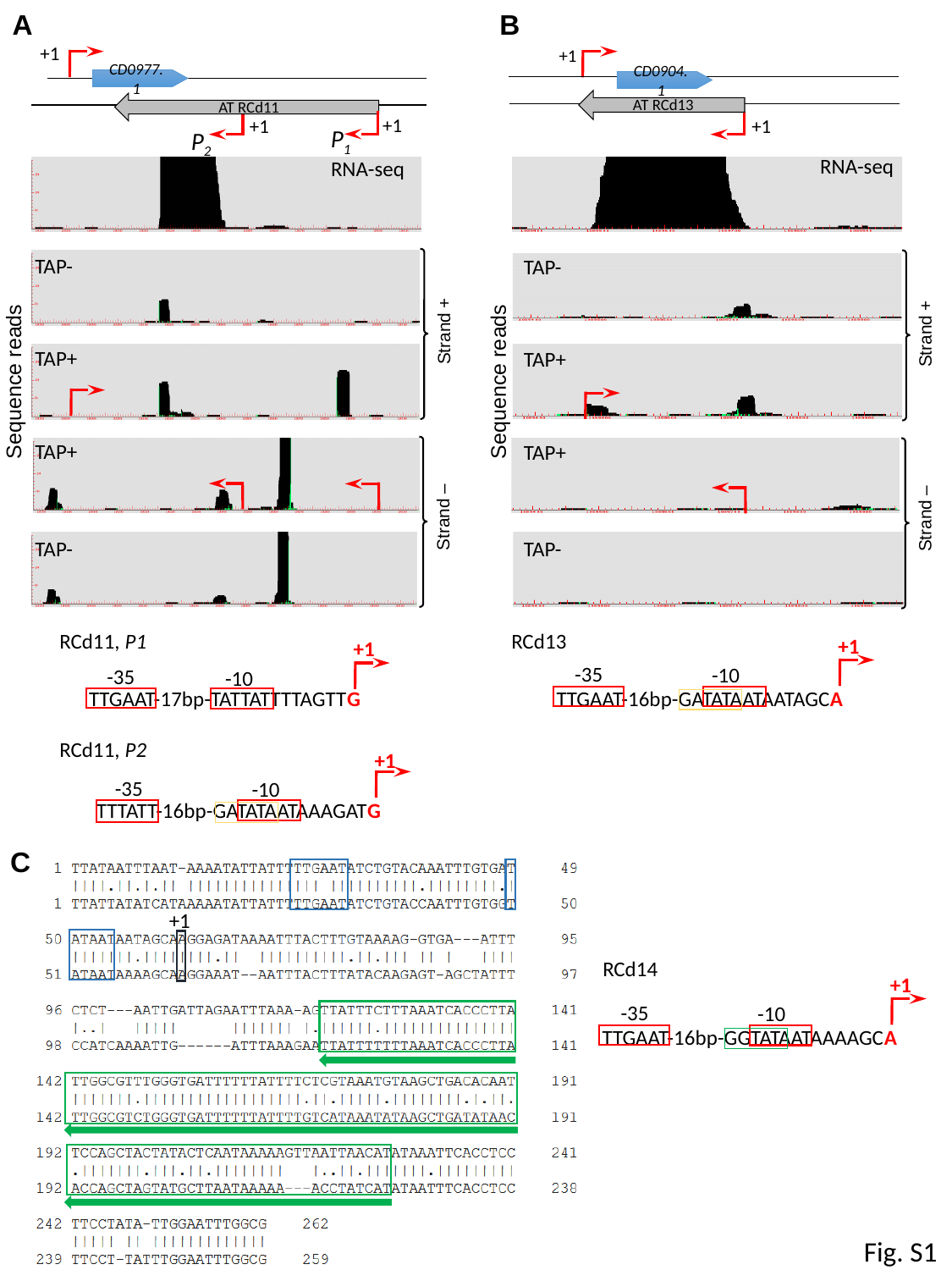

B
A
+1
+1
CD0977.1
CD0904.1
AT RCd13
AT RCd11
+1
+1
+1
P_1_
P_2_
RNA-seq
RNA-seq
TAP-
TAP-
Strand +
Strand +
TAP+
TAP+
Sequence reads
Sequence reads
TAP+
TAP+
Strand –
Strand –
TAP-
TAP-
RCd11, P1
+1
-35
-10
TTGAAT-17bp-TATTATTTTAGTTG
RCd13
+1
-35
-10
TTGAAT-16bp-GATATAATAATAGCA
RCd11, P2
+1
-35
-10
TTTATT-16bp-GATATAATAAAGATG
C
0
+1
RCd14
+1
-35
-10
TTGAAT-16bp-GGTATAATAAAAGCA
Fig. S1

### Slide 2
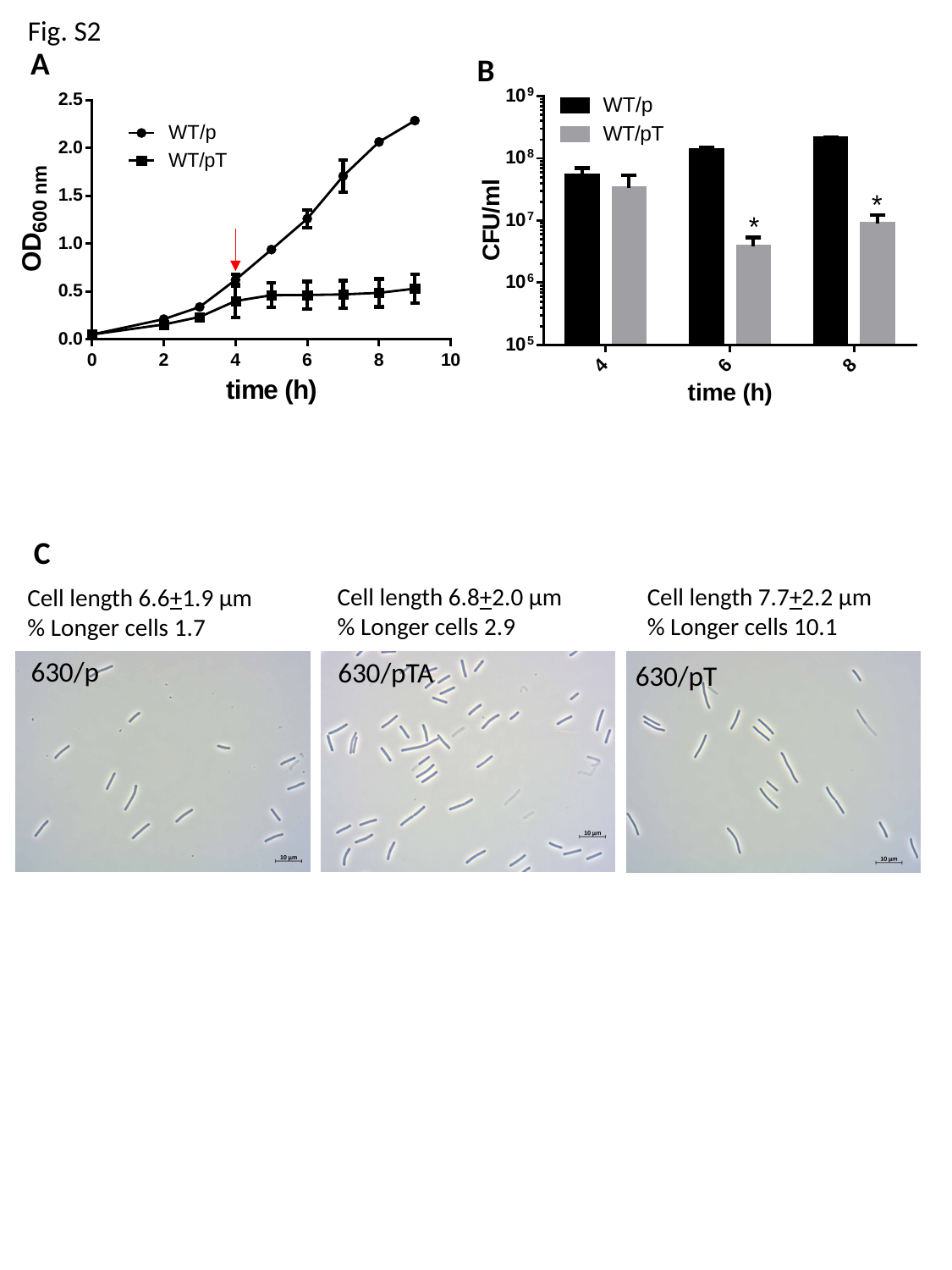

Fig. S2
A
B
C
Cell length 6.8+2.0 µm
% Longer cells 2.9
Cell length 7.7+2.2 µm
% Longer cells 10.1
Cell length 6.6+1.9 µm
% Longer cells 1.7
630/p
630/pTA
630/pT

### Slide 3
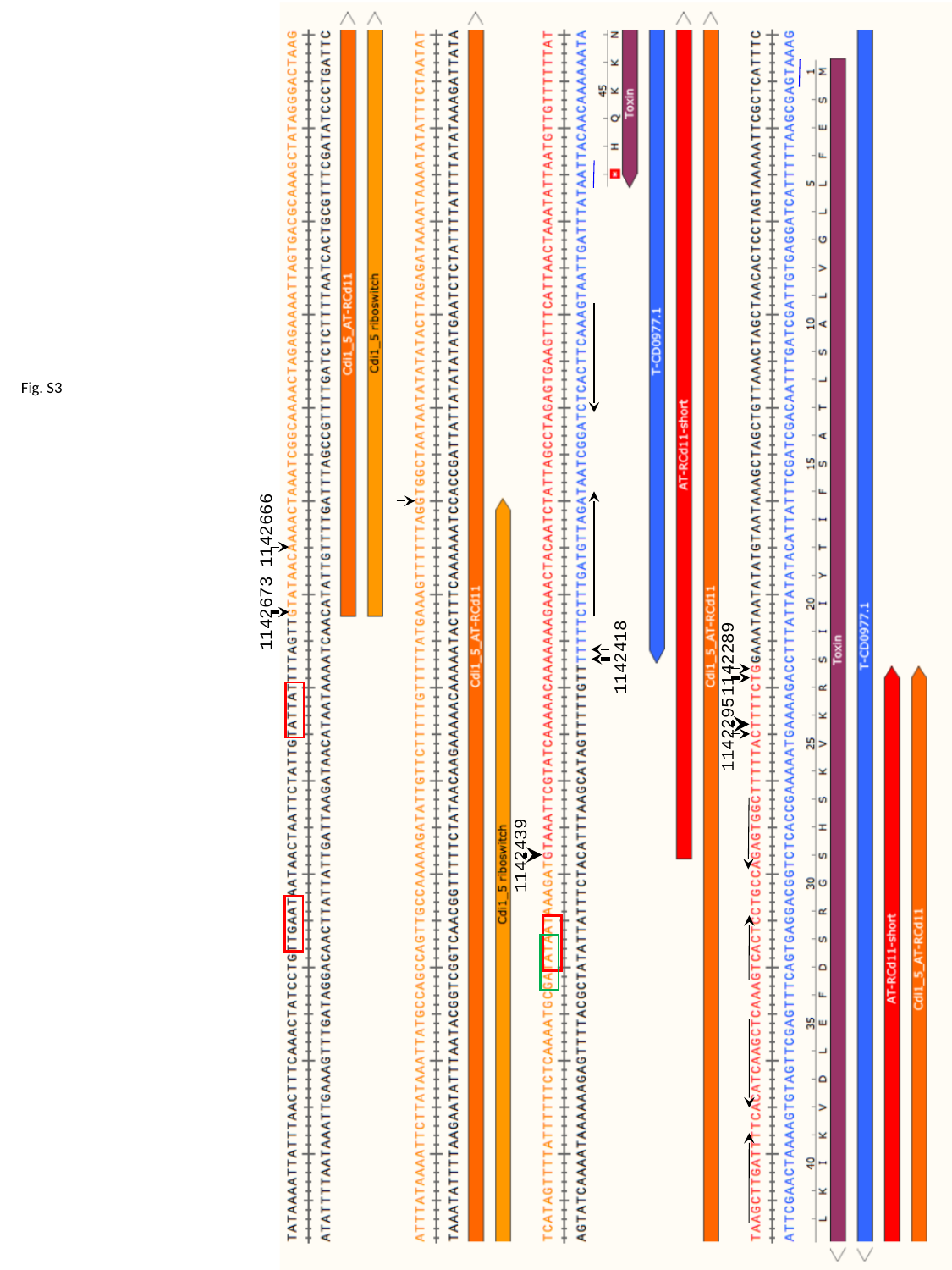

1142666
1142673
1142439
1142418
1142289
1142295
1142137
1142158
Fig. S3

### Slide 4
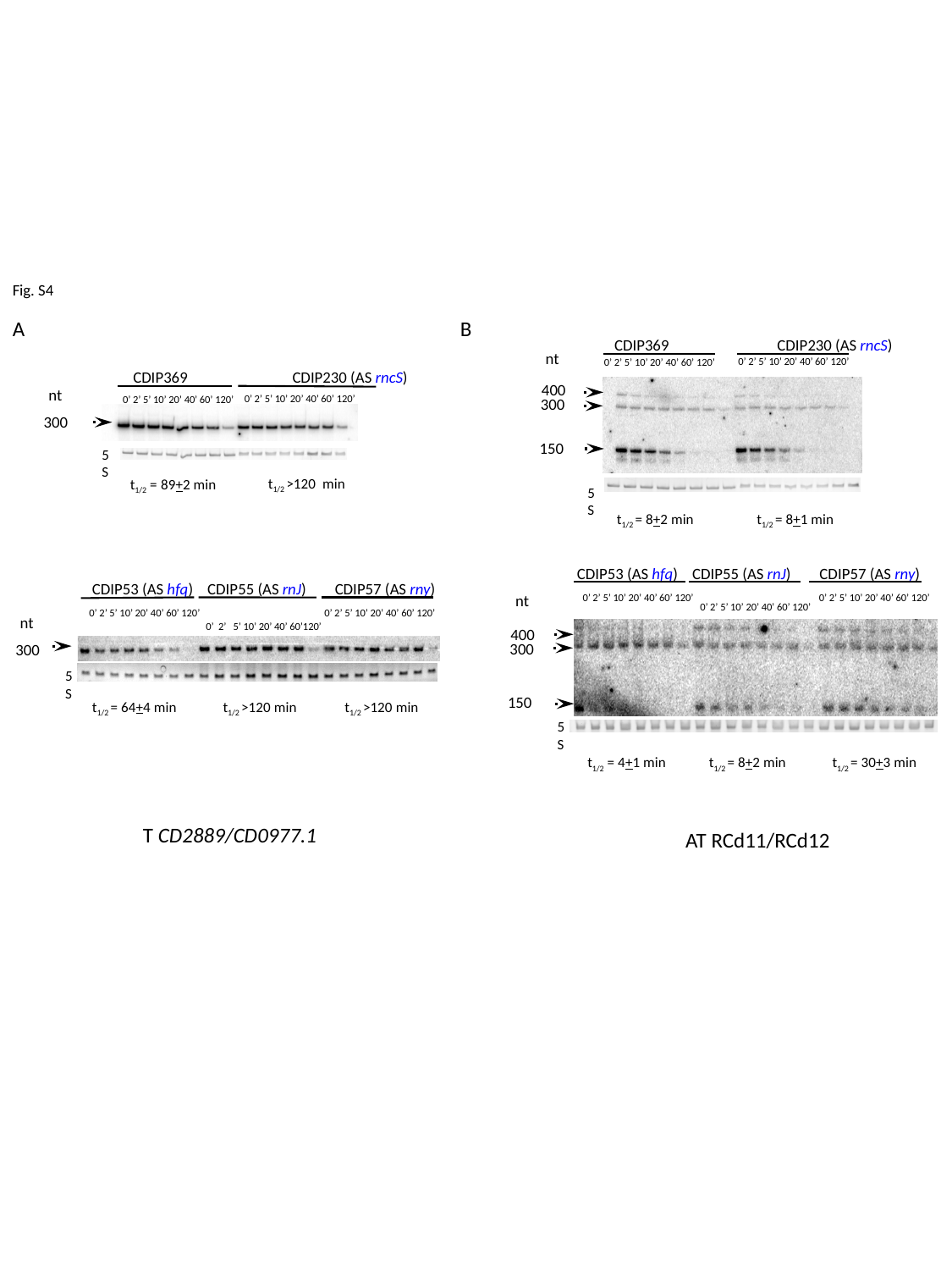

Fig. S4
A
B
CDIP369	 CDIP230 (AS rncS)
nt
0’ 2’ 5’ 10’ 20’ 40’ 60’ 120’
0’ 2’ 5’ 10’ 20’ 40’ 60’ 120’
CDIP369	 CDIP230 (AS rncS)
400
nt
0’ 2’ 5’ 10’ 20’ 40’ 60’ 120’
0’ 2’ 5’ 10’ 20’ 40’ 60’ 120’
300
300
150
5S
t_1/2_ >120 min
t_1/2_ = 89+2 min
5S
t_1/2_ = 8+2 min
t_1/2_ = 8+1 min
CDIP53 (AS hfq) CDIP55 (AS rnJ) CDIP57 (AS rny)
CDIP53 (AS hfq) CDIP55 (AS rnJ) CDIP57 (AS rny)
0’ 2’ 5’ 10’ 20’ 40’ 60’ 120’
nt
0’ 2’ 5’ 10’ 20’ 40’ 60’ 120’
0’ 2’ 5’ 10’ 20’ 40’ 60’ 120’
0’ 2’ 5’ 10’ 20’ 40’ 60’ 120’
0’ 2’ 5’ 10’ 20’ 40’ 60’ 120’
nt
0’ 2’ 5’ 10’ 20’ 40’ 60’120’
400
300
300
5S
150
t_1/2_ = 64+4 min
t_1/2_ >120 min
t_1/2_ >120 min
5S
t_1/2_ = 4+1 min
t_1/2_ = 8+2 min
t_1/2_ = 30+3 min
T CD2889/CD0977.1
AT RCd11/RCd12

### Slide 5
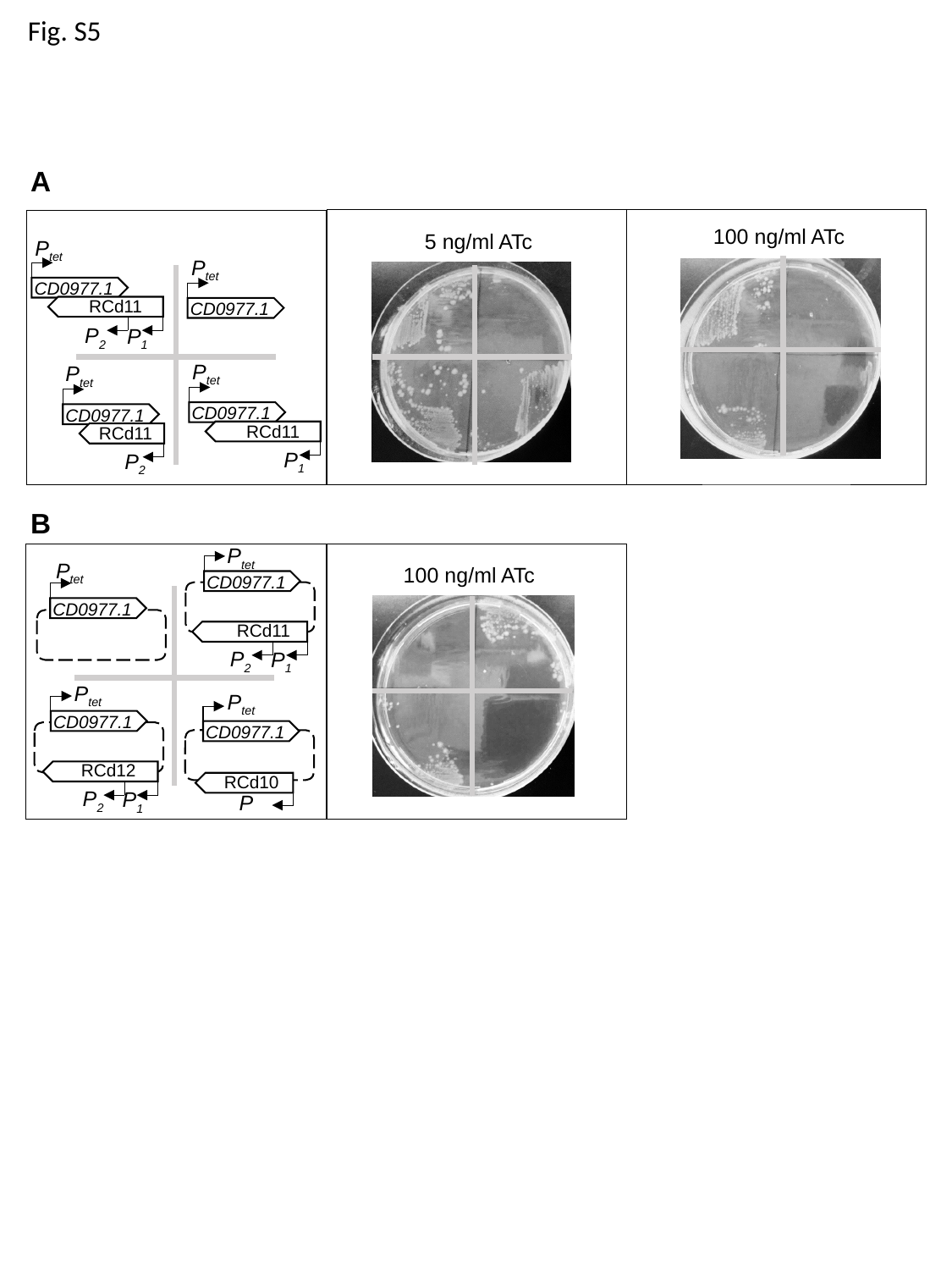

Fig. S5
A
100 ng/ml ATc
5 ng/ml ATc
P_tet_
CD0977.1
RCd11
P_2_
P_1_
P_tet_
CD0977.1
P_tet_
CD0977.1
RCd11
P_1_
P_tet_
CD0977.1
RCd11
P_2_
B
P_tet_
CD0977.1
RCd11
P_2_
P_1_
P_tet_
CD0977.1
100 ng/ml ATc
P_tet_
CD0977.1
RCd12
P_2_
P_1_
P_tet_
CD0977.1
RCd10
P

### Slide 6
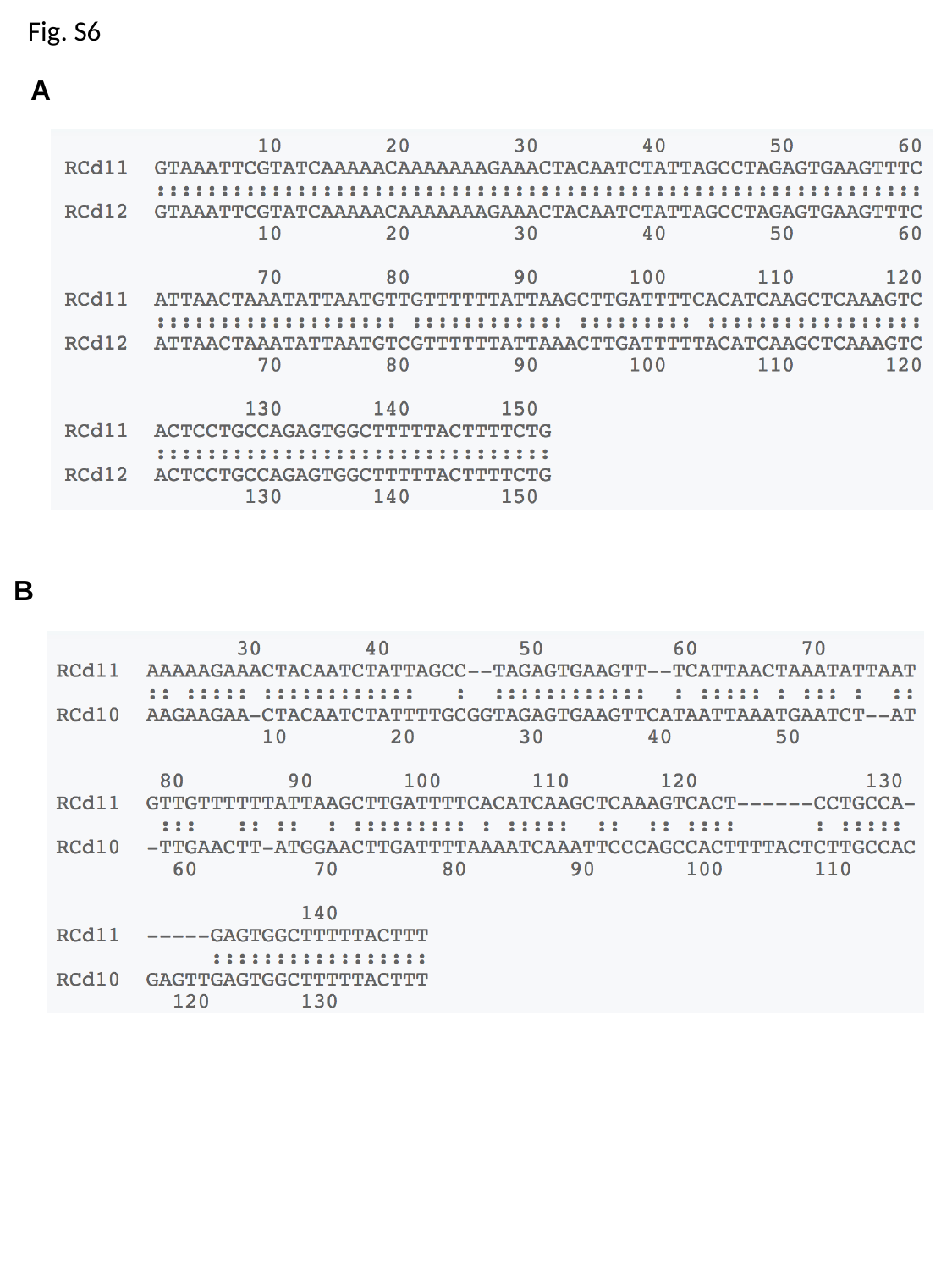

Fig. S6
A
B

### Slide 7
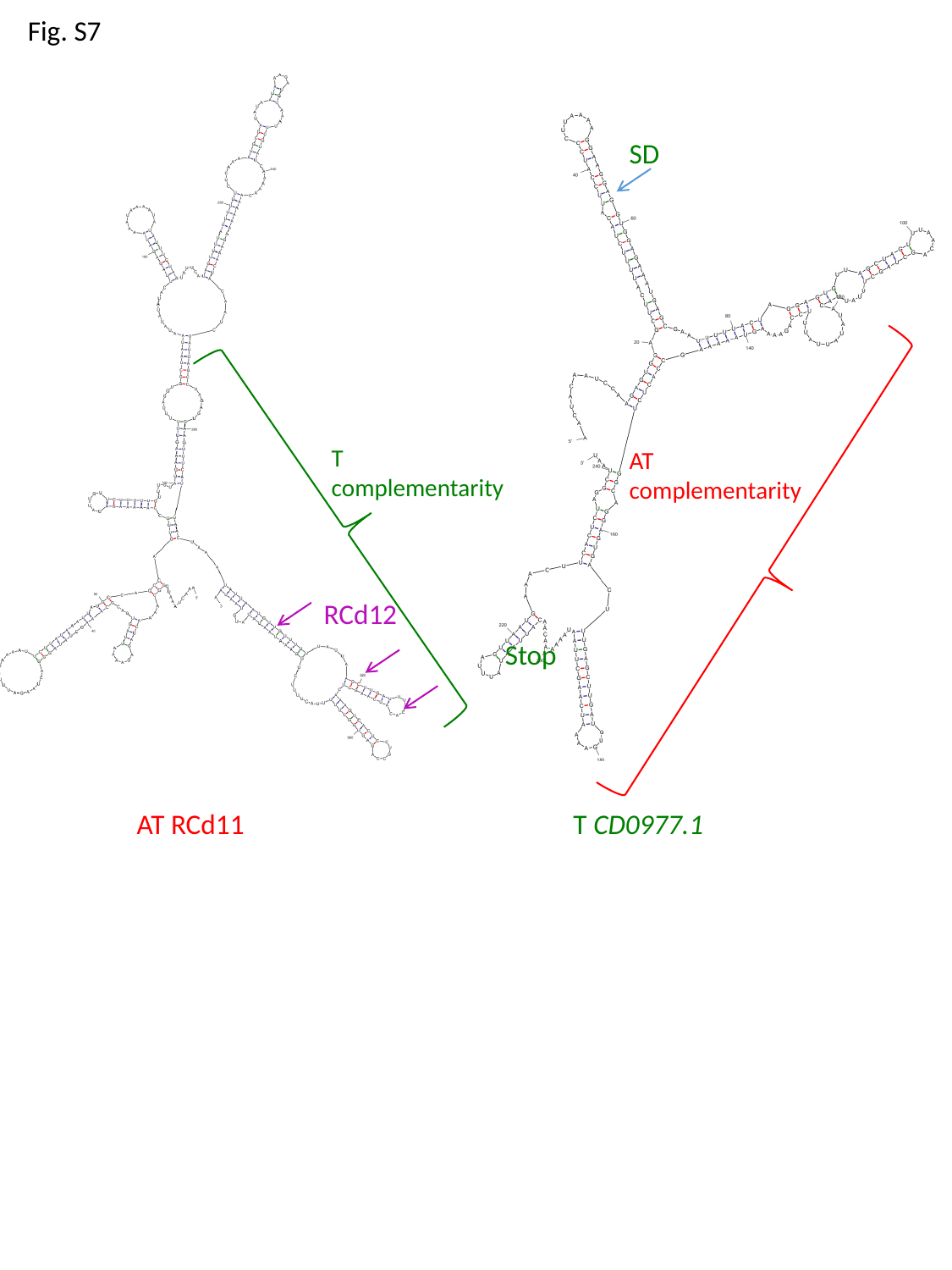

Fig. S7
SD
AT complementarity
Stop
T complementarity
RCd12
AT RCd11
T CD0977.1

### Slide 8
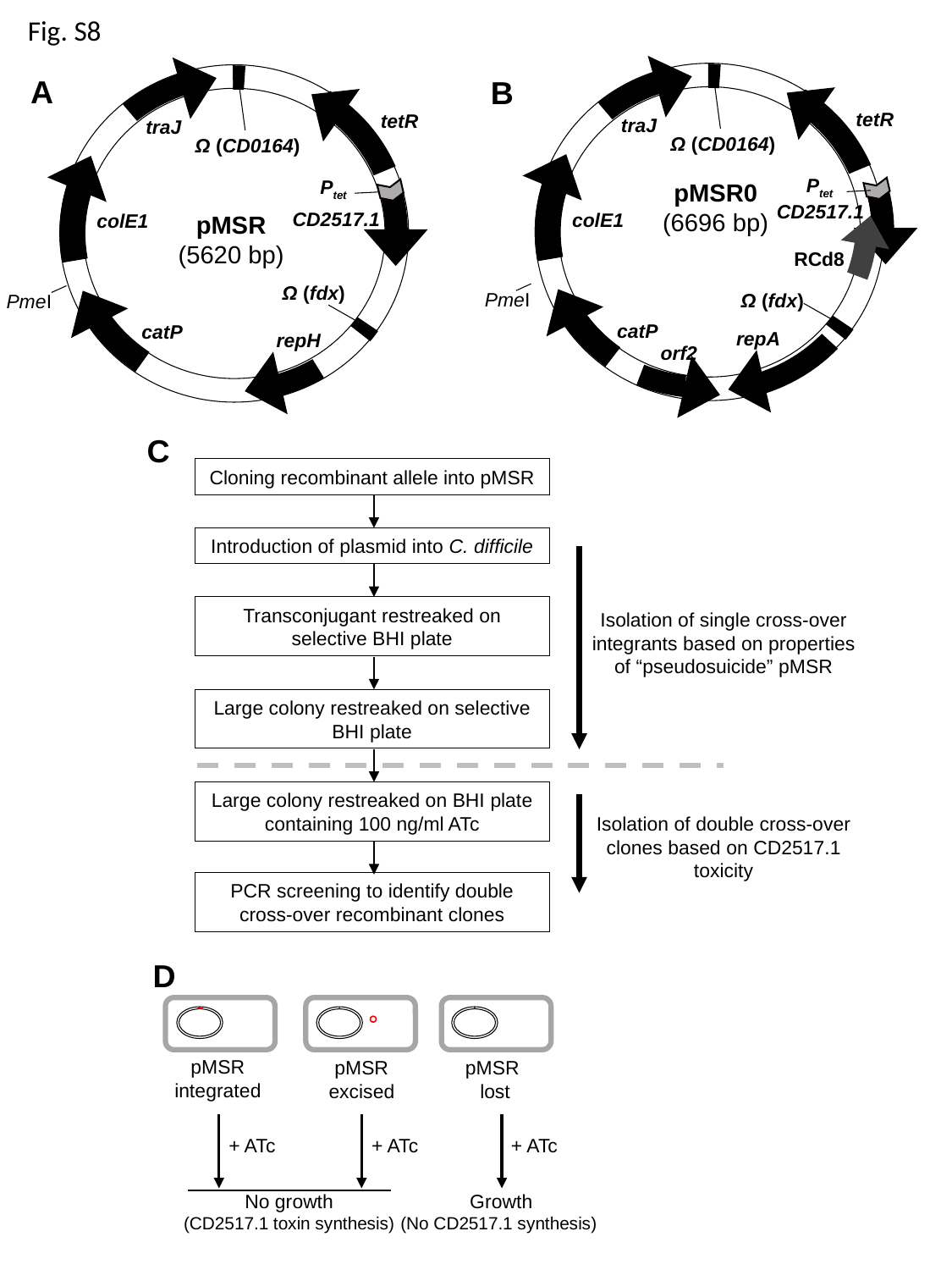

Fig. S8
tetR
traJ
Ω (CD0164)
P_tet_
pMSR0
(6696 bp)
colE1
RCd8
Ω (fdx)
catP
repA
A
B
tetR
traJ
Ω (CD0164)
P_tet_
CD2517.1
CD2517.1
colE1
pMSR
(5620 bp)
Ω (fdx)
PmeI
PmeI
catP
repH
orf2
C
Cloning recombinant allele into pMSR
Introduction of plasmid into C. difficile
Transconjugant restreaked on selective BHI plate
Isolation of single cross-over integrants based on properties of “pseudosuicide” pMSR
Large colony restreaked on selective BHI plate
Large colony restreaked on BHI plate containing 100 ng/ml ATc
Isolation of double cross-over clones based on CD2517.1 toxicity
PCR screening to identify double cross-over recombinant clones
D
pMSR integrated
pMSR excised
pMSR
lost
+ ATc
+ ATc
+ ATc
No growth
(CD2517.1 toxin synthesis)
Growth
(No CD2517.1 synthesis)

### Slide 9
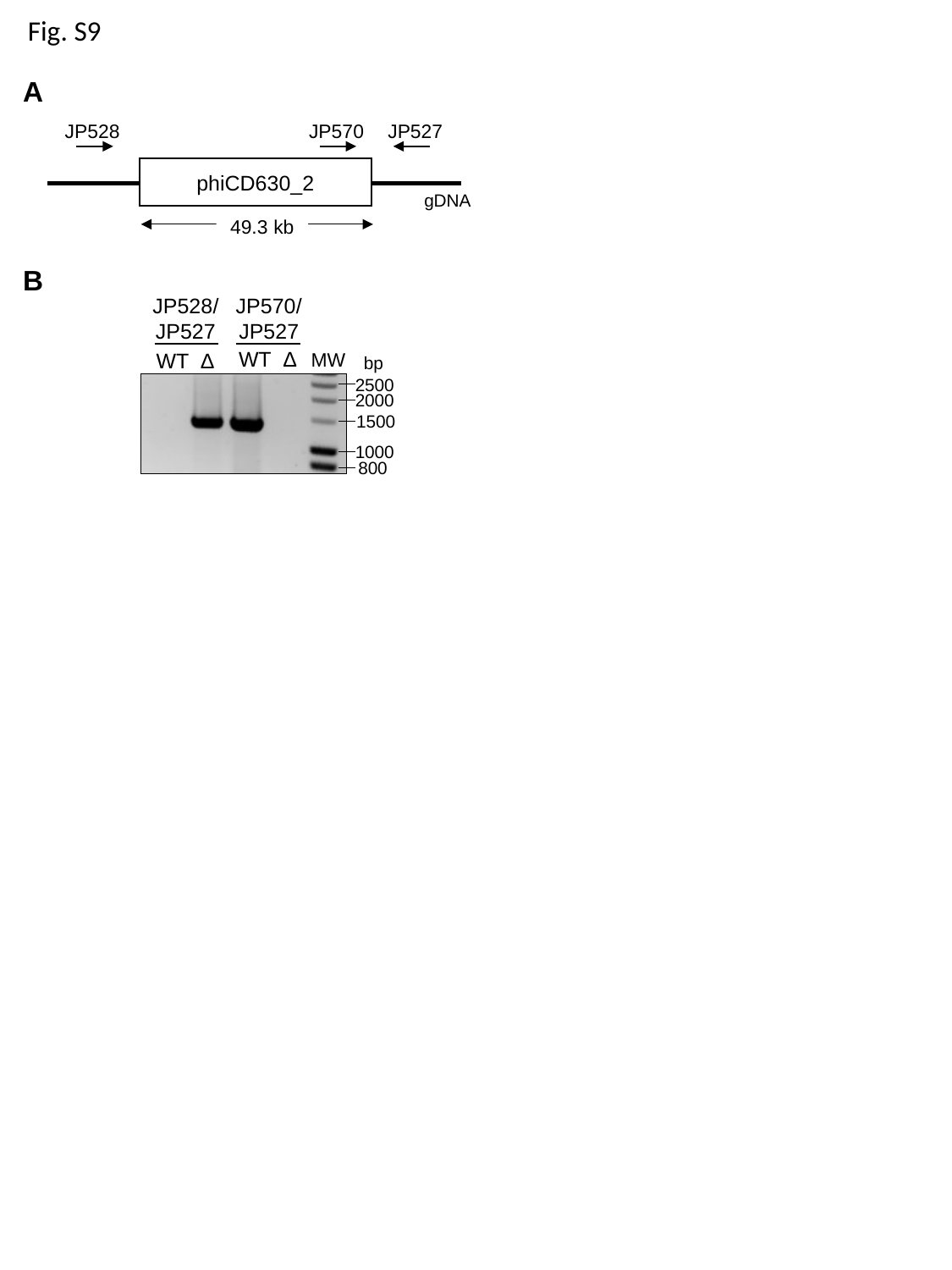

Fig. S9
A
JP528
JP570
JP527
phiCD630_2
gDNA
49.3 kb
B
JP528/JP527
JP570/JP527
WT Δ
WT Δ
MW
bp
2500
2000
1500
1000
800

### Slide 10
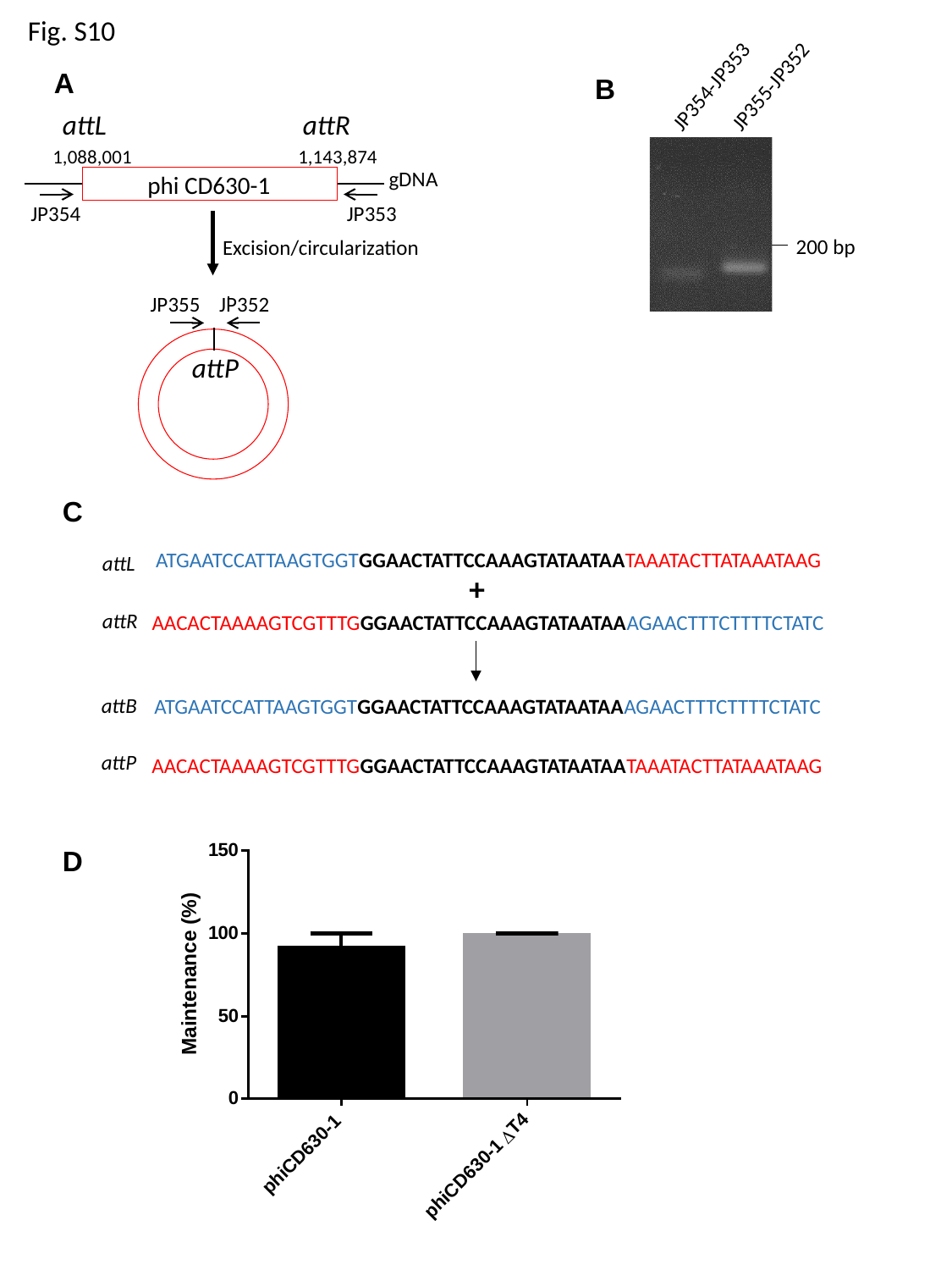

Fig. S10
JP355-JP352
JP354-JP353
A
B
attL
attR
1,088,001
1,143,874
gDNA
phi CD630-1
JP354
JP353
200 bp
Excision/circularization
JP355
JP352
attP
C
ATGAATCCATTAAGTGGTGGAACTATTCCAAAGTATAATAATAAATACTTATAAATAAG
attL
+
attR
AACACTAAAAGTCGTTTGGGAACTATTCCAAAGTATAATAAAGAACTTTCTTTTCTATC
attB
ATGAATCCATTAAGTGGTGGAACTATTCCAAAGTATAATAAAGAACTTTCTTTTCTATC
attP
AACACTAAAAGTCGTTTGGGAACTATTCCAAAGTATAATAATAAATACTTATAAATAAG
D

### Slide 11
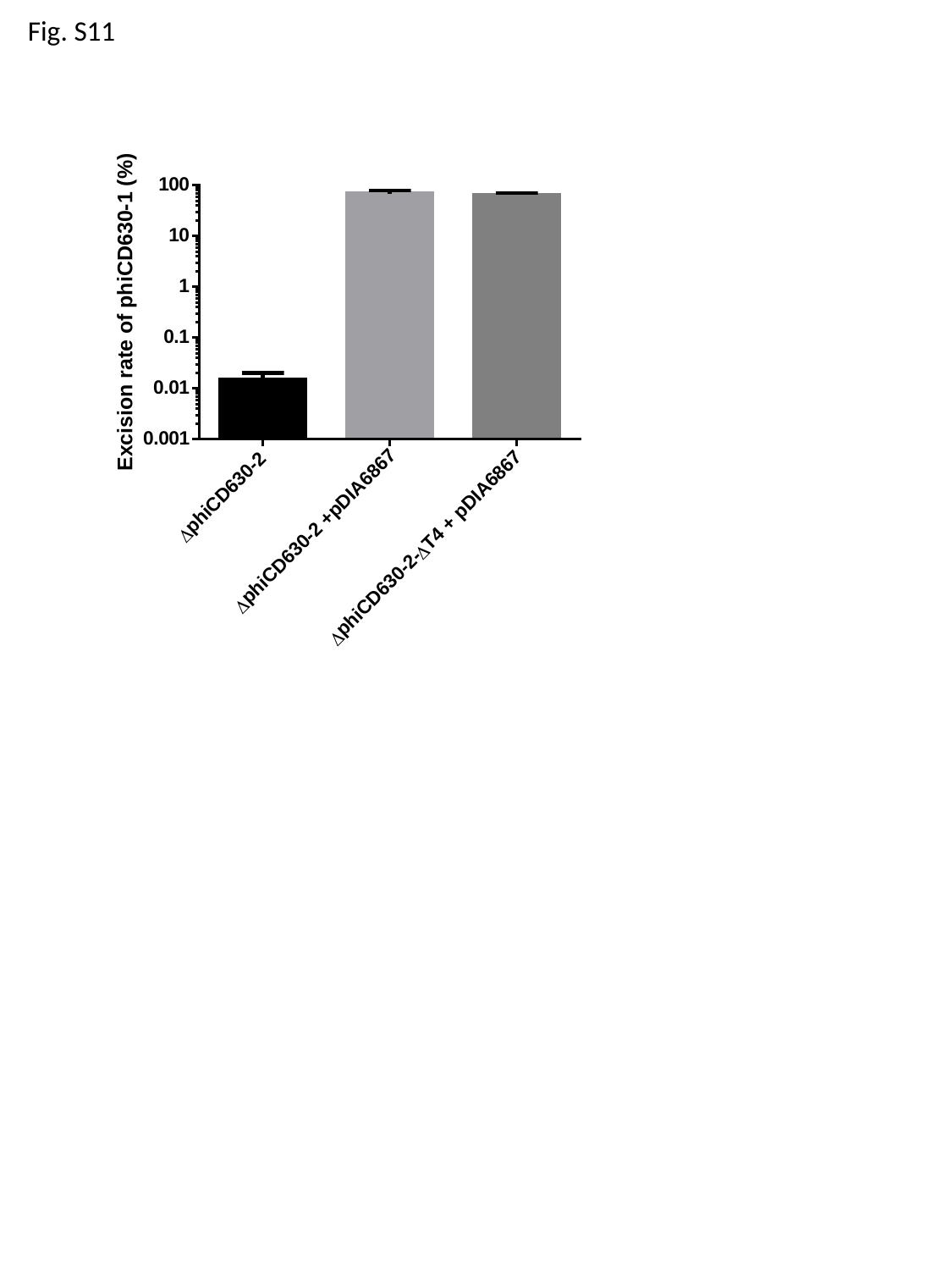

Fig. S11

### Slide 12
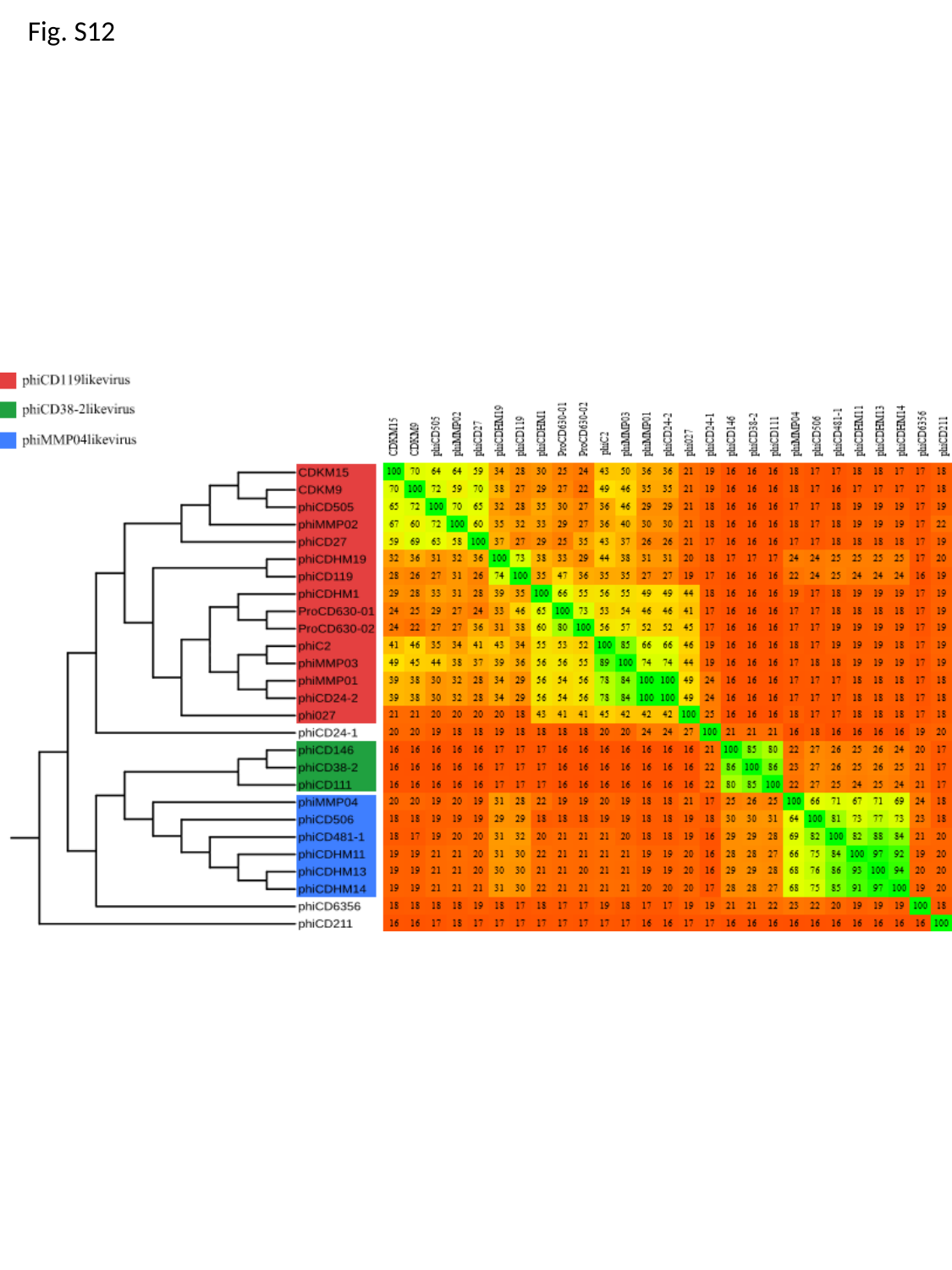

Fig. S12

### Slide 13
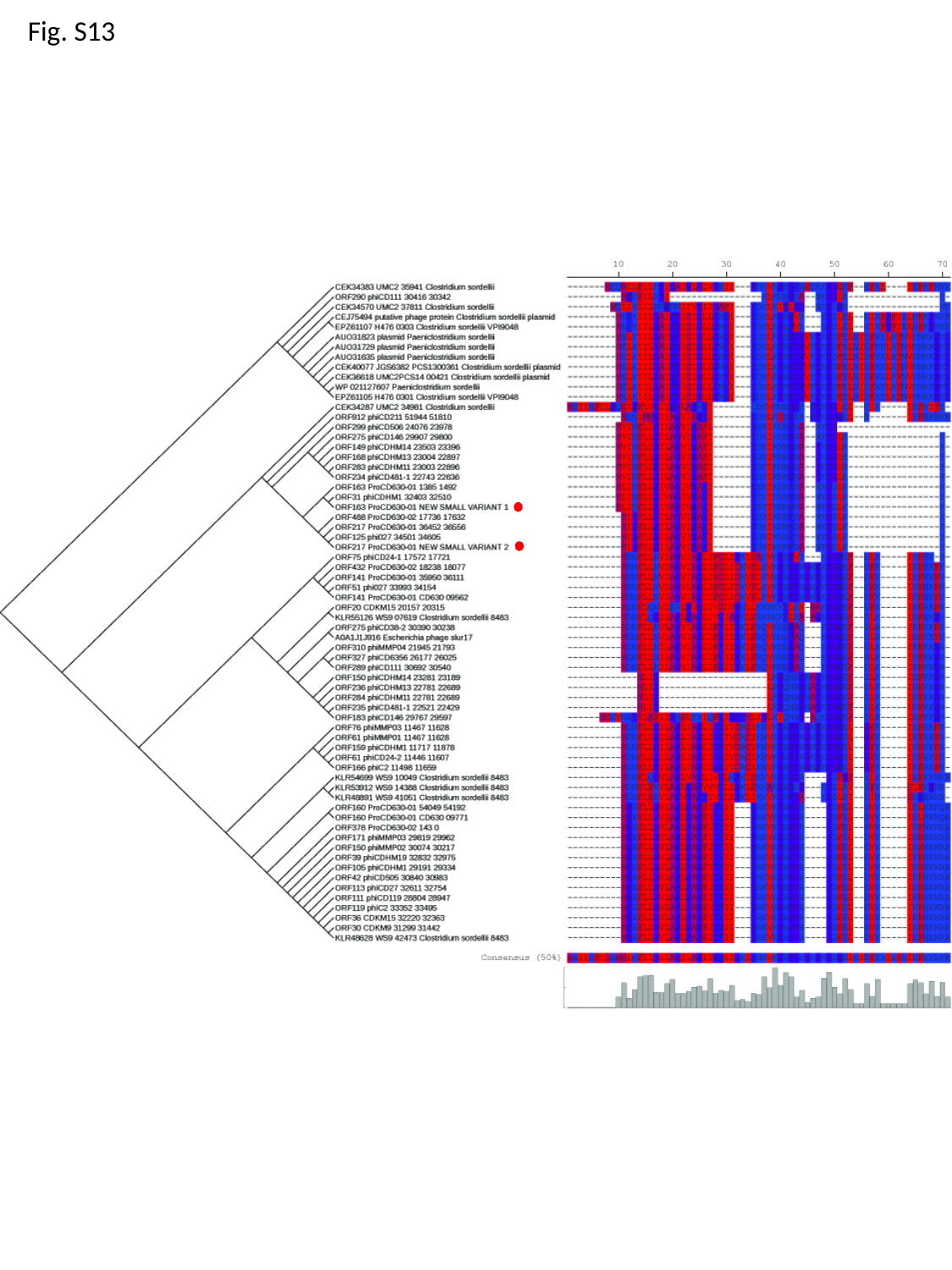

Fig. S13
