## Supplemental Data 1 for "Type I toxin-antitoxin systems contribute to mobile genetic elements maintenance in *Clostridioides difficile* and can be used as a counter-selectable marker for chromosomal manipulation"

**Table S1. Strains and plasmids used in this study.**

| **Strain** | **Genotype** | **Origin** |
| --- | --- | --- |
| ***E. coli*** |  |  |
| NEB-10 beta | Δ*(ara-leu) 7697 araD139  fhuA* Δ*lacX74 galK16 galE15 e14-* ϕ*80*d*lacZ*Δ*M15  recA1 relA1 endA1 nupG  rpsL* (Str^R^) *rph spoT1* Δ*(mrr-hsdRMS-mcrBC)* | New England Biolabs |
| DH5α | F‑ Φ80*lac*ZΔM15 Δ(*lacZYA‑argF*) U169 *recA1* *endA1* *hsdR17* (rK‑, mK+) *phoA* *supE*44 λ‑ *thi*‑1 *gyrA*96 *relA*1 | Invitrogen |
| HB101 (RP4) | *supE44* *aa14 galK2 lacY1* Δ(*gpt-proA*) 62 *rpsL20 (*Str^R^*)xyl-5* *mtl-1 recA13* Δ(*mcrC-mrr*) *hsdS*_B_ (r_B_-m_B_-) RP4 (Tra^+^ IncP Ap^R^ Km^R^ Tc^R^) | Laboratory stock |
| ***C. difficile*** |  |  |
| 630Δ*erm* | 630 Δ*ermB* | Laboratory stock (1) |
| CDIP51 | 630 Δ*erm* strain carrying pRPF185 vector | (2) |
| CDIP369 (630/p) | 630 Δ*erm* strain carrying pRPF185Δ*gusA* vector | pDIA6103🡪630Δ*erm* |
| CDIP0506 | 630 Δ*erm* strain carrying pDIA6335 plasmid for inducible expression of *CD0977.1* | pDIA6335🡪630Δ*erm* |
| CDIP0966 | 630 Δ*erm* Δ*CD0977.1* | Δ*CD0977.1*🡪630Δ*erm* |
| CDIP0997 | 630 Δ*erm* strain carrying pDIA6622 plasmid for inducible expression of HA-tagged *CD0977.1* | pDIA6622🡪630Δ*erm* |
| CDIP1174 | 630 Δ*erm* Δ*CD0956.2* Δ*CD0977.1* | Δ*CD0956.2*🡪CDIP0966 |
| CDIP1266 | 630 Δ*erm* strain carrying pDIA6785 plasmid for inducible expression of *CD0977.1* and co-expression of RCd11 from its two own promoters | pDIA6785🡪630Δ*erm* |
| CDIP1268 | 630 Δ*erm* strain carrying pDIA6787 plasmid for expression of *CD0977.1* and co-expression of RCd11 from their native promoters | pDIA6787🡪630Δ*erm* |
| CDIP1270 | 630 Δ*erm* strain carrying pDIA6791 plasmid for inducible expression of *CD0977.1* and expression of RCd11 from its two own promoters in *trans* | pDIA6791🡪630Δ*erm* |
| CDIP1271 | 630 Δ*erm* strain carrying pDIA6792 plasmid for inducible expression of *CD0977.1* and expression of RCd12 from its two own promoters in *trans* | pDIA6792🡪630Δ*erm* |
| CDIP1272 | 630 Δ*erm* strain carrying pDIA6793 plasmid for inducible expression of *CD0977.1* and expression of RCd8 in *trans* | pDIA6793🡪630Δ*erm* |
| CDIP1338 | 630 Δ*erm* ΔPhiCD630_2 | ΔPhiCD630_2🡪630Δ*erm* |
| CDIP1339 | 630 Δ*erm* Δ*CD0956.2* Δ*CD0977.1* ΔPhiCD630_2 | ΔPhiCD630_2🡪CDIP1174 |
| CDIP1368 | 630 Δ*erm* strain carrying pDIA6934 plasmid for inducible expression of *CD0904.1* | pDIA6866🡪630Δ*erm* |
| CDIP1373 | 630 Δ*erm* Δ*CD0904.1*Δ*CD0956.2* Δ*CD0977.1* ΔPhiCD630_2 | Δ*CD0904.1*🡪CDIP1339 |
| CDIP1374 | 630 Δ*erm* Δ*CD0904.1* Δ*CD0956.2* Δ*CD0956.2* Δ*CD0977.1* ΔPhiCD630_2 | Δ*CD0956.3*🡪CDIP1373 |
| CDIP1434 | 630 Δ*erm* strain carrying pDIA6934 plasmid for inducible expression of *CD0904.1* and co-expression of RCd13 | pDIA6934🡪630Δ*erm* |
| CDIP1440 | 630 Δ*erm* strain carrying pDIA6947 plasmid for expression of *CD0904.1* and co-expression of RCd13 from their native promoters | pDIA6947🡪630Δ*erm* |
| CDIP1441 | 630 Δ*erm* strain carrying pDIA6948 plasmid for expression of *CD0956.2* and co-expression of RCd10 from their native promoters | pDIA6948🡪630Δ*erm* |
| CDIP1442 | 630 Δ*erm* strain carrying pDIA6949 plasmid for expression of *CD0956.3* and co-expression of RCd14 from their native promoters | pDIA6949🡪630Δ*erm* |
| CDIP1510 | 630 Δ*erm* ΔPhiCD630_2 phiCD630_1::P*_thl_*-*ermB* | P*_thl_*-*ermB*🡪CDIP1338 |
| CDIP1511 | 630 Δ*erm* ΔPhiCD630_2 phiCD630_1::P*_thl_*-*ermB*-ΔT4 | P*_thl_*-*ermB*🡪CDIP1374 |
| CDIP1519 | 630 Δ*erm* ΔPhiCD630_2 phiCD630_1::P*_thl_*-*ermB* carrying pDIA6867 for inducible expression of *CD0912* | pDIA6867🡪CDIP1510 |
| CDIP1520 | 630 Δ*erm* ΔPhiCD630_2 phiCD630_1::P*_thl_*-*ermB*-ΔT4 carrying pDIA6867 for inducible expression of *CD0912* | pDIA6867🡪CDIP1511 |
| CDIP1593 | 630 Δ*erm* strain carrying pDIA7030 plasmid for inducible expression of *CD0956.3* | pDIA7030🡪630Δ*erm* |
| CDIP1594 | 630 Δ*erm* strain carrying pDIA7031 plasmid for inducible expression of *CD0956.3* and co-expression of RCd14 | pDIA7031🡪630Δ*erm* |
| **Plasmid** |  |  |
| pMTL84121 | Tm^R,^ expression and cloning vector | (3) |
| pRPF185 | *P_tet_-gusA* Tm^R,^ expression and cloning vector | (4) |
| pMSR | Allele exchange in *C. difficile* 630 | This work |
| pMSR0 | Allele exchange in *C. difficile* R20291 | This work |
| pDIA6103 | pRPF185 Δ*gus* vector derivative | (5) |
| pDIA6319 | pRPF185 derivative carrying *P_tet_*-*CD2517.1* for inducible *CD2517.1* toxin expression | (6) |
| pDIA6335 | pRPF185 derivative carrying *P_tet_*-*CD0977.1* for inducible toxin expression | This work |
| pDIA6507 | pMSR derivative for *CD09956.2* deletion | This work |
| pDIA6622 | pRPF185 derivative carrying *P_tet_*-*CD0977.1* for inducible *CD0977.1*-HA-tag toxin expression | This work |
| pDIA6755 | pMSR derivative for *CD0977.1* deletion | This work |
| pDIA6785 | pRPF185 derivative carrying *P_tet_*-*CD0977.1-*RCd11*-P_2_-P_1_* for inducible toxin expression and co-expression of AT with both promoters | This work |
| pDIA6787 | pMTL84121 derivative carrying *CD0977.1* and its cognate antitoxin with their respective native promoters | This work |
| pDIA6791 | pRPF185 derivative carrying *Ptet*-  *CD0977.1* and *P_1_-P_2_-*RCd11 on a second plasmid site for inducible *CD0977.1* toxin expression and expression of RCd11 in *trans* | This work |
| pDIA6792 | pRPF185 derivative carrying *Ptet*-  *CD0977.1* and *P_1_-P_2_-*RCd12 on a second plasmid site for inducible *CD0977.1* toxin expression and expression of RCd12 in *trans* | This work |
| pDIA6793 | pRPF185 derivative carrying *Ptet*-  *CD0977.1* and *P-RCd10* on a second plasmid site for inducible *CD0977.1* toxin expression and expression of RCd10 in *trans* | This work |
| pDIA6816 | pRPF185 derivative carrying *P_tet_*-*CD0977.1-*RCd11*-P_2_* for inducible toxin expression and co-expresssion of RCd11 with second promoter only | This work |
| pDIA6817 | pRPF185 derivative carrying *P_tet_*-*CD0977.1-*RCd11*-P_1_* for inducible toxin expression and co-expresssion of RCd11 with first promoter only | This work |
| pDIA6835 | pMSR derivative for PhiCD630_2 deletion | This work |
| pDIA6836 | pMSR derivative for PhiCD630_1 deletion | This work |
| pDIA6857 | pMSR derivative for *CD0956.3* deletion | This work |
| pDIA6858 | pMSR derivative for *CD0904.1* deletion | This work |
| pDIA6866 | pRPF185 derivative carrying *P_tet_*-*CD0904.1* | This work |
| pDIA6867 | pRPF185 derivative carrying *P_tet_*-*CD0912* | This work |
| pDIA6934 | pRPF185 derivative carrying *P_tet_*-*CD0904.1* and co-expression of RCd13 | This work |
| pDIA6947 | pMTL84121 derivative carrying *CD0904.1* and its cognate antitoxin with their respective native promoters | This work |
| pDIA6948 | pMTL84121 derivative carrying *CD0956.2* and its cognate antitoxin with their respective native promoters | This work |
| pDIA6949 | pMTL84121 derivative carrying *CD0956.3*and its cognate antitoxin with their respective native promoters | This work |
| pDIA6976 | pMSR derivative for *Pthl-ermB* insertion within PhiCD630-1 | This work |
| pDIA7030 | pRPF185 derivative carrying *P_tet_*-*CD0956.3* | This work |
| pDIA7031 | pRPF185 derivative carrying *P_tet_*-*CD0956.3* and co-expression of RCd14 | This work |
