## Supplemental Data 2 for "Type I toxin-antitoxin systems contribute to mobile genetic elements maintenance in *Clostridioides difficile* and can be used as a counter-selectable marker for chromosomal manipulation"

**Table S2. Oligonucleotides used in this study.**

| **Name** | **Sequence (5'-3')** | **Description** |
| --- | --- | --- |
| **pDIA6103 cloning** | |  |
| IMV507 | GGGATTTCTCACATAAAATAGAG | 5’pDIA6103 insert screening |
| IMV508 | TAAAATAAGCTTGATCGTAGCG | 3’pDIA6103 insert screening |
| JP420 | GGATCCTATAAGTTTTAATAAAACTTTAAATAG | 5’ pDIA6103 linearization for *P_tet_* cloning with Hifi DNA assembly |
| JP421 | AGGCCTGGAGCTCAGA | 3’ pDIA6103 linearization for *P_tet_* cloning with Hifi DNA assembly |
| JP473 | aacatctgagctccaggcctAACTACAATCCAAGAGTG | *5’CD0977.1-*RCd11*-*  *P_2_-P_1_* Hifi DNA assembly |
| JP474 | tattaaaacttataggatccGGAGCTTTTTAGGACAAAAAC | *3’CD0977.1-*RCd11*-*  *P_2_-P_1_* Hifi DNA assembly |
| JP492 | tattaaaacttataggatccGGTGGCTAATAATATATATACTTAGAG | *3’CD0977.1-*RCd11*-*  *P_2_*  Hifi DNA assembly |
| JP574 | tattaaaacttataggatccGAATTATTTTTTTAAATCACCCTTATTG | *3’CD0956.3* Hifi DNA assembly |
| JP592 | cagatctgagctccaggcctGACGAACTAGTTGGAAGATAG | *5’CD0912* Hifi DNA assembly |
| JP593 | tattaaaacttataggatccCAACTAAATCACCCCCTTTC | *3’CD0912* Hifi DNA assembly |
| JP594 | cagatctgagctccaggcctCGTATAAACGCCAAATTC | *5’CD0904.1* Hifi DNA assembly |
| JP595 | tattaaaacttataggatccGATTAGAATTTAAAAGTTATTTCTTTAAATC | *3’CD0904.1* Hifi DNA assembly |
| JP635 | tattaaaacttataggatccCAATTATAATTTAATAAAATATTATTTTTGAATATCTG | *3’CD0904.1*-RCd13-*P* Hifi DNA assembly |
| JP694 | cagatctgagctccaggcctcgccaaattccaaataag | *5’CD0956.3* Hifi DNA assembly |
| JP695 | tattaaaacttataggatccgctcactgcaaaatctctac | *3’CD0956.3*-RCd14-*P* Hifi DNA assembly |
| OS650 | GAAGGCCTAACTACAATCCAAGAGTGGAGCTTCATTTTC | *5’CD0977.1-*StuI |
| OS651 | GGGGATCCAAAGATGTAAATTCGTATCAAAAAC | *3’CD0977.1-*BamHI |
| OS652 | GAAGGCCTTAACATATTTAGTATATACCTATG | *5’CD0977.1*-RCd11- *P_2_-P_1_* -StuI |
| OS653 | GGGGATCCGGAGCTTTTTAGGACAAAAACTAT | *5’CD0977.1*-RCd11- *P_2_-P_1_* -BamHI |
| **pDIA6103 second cloning site (distant from the MCS)** | |  |
| JP363 | CAGATTACGCTTAATATTTAGTTAATGAAACTTCACTCTAGG | 3’ HA-tagging on the 3' extremity of CD2517.1 |
| JP326 | GAACATCGTATGGGTAATGTTGTTTTTTATTAAGCTTGATTTTC | 5’’ HA-tagging on the 3' extremity of CD2517.1 |
| JP403 | CTGTATCGTAACTAGAGAACCAAAC | 5’ plasmid linearization for cloning into a site different from the MCS with Hifi DNA assembly |
| JP404 | GGGATATGCTTATATTGAGTTATAGTAC | 5’ plasmid linearization for cloning into a site different from the MCS with Hifi DNA assembly |
| JP409 | CTAACATACATCATAGTTACTAAACTATGG | 5’pRPF185-screening |
| JP410 | GCTTTTTCCGTCGTTTG | 3’pRPF185-screening |
| JP476 | actcaatataagcatatcccGGAGCTTTTTAGGACAAAAAC | 5’ *P_1_-P_2_-*RCd11 Hifi DNA assembly |
| JP477 | gttctctagttacgatacagCCAGAAAAGTAAAAAGCC | 3’ *P_1_-P_2_-*RCd11 Hifi DNA assembly |
| JP478 | actcaatataagcatatcccCATTTTTTTTATATAAACAATGAAATTCAAG | 5’ *P-RCd8* Hifi DNA assembly |
| JP479 | gttctctagttacgatacagCTACAATTTATAGAGTGGAGTTC | 3’ *P-RCd8* Hifi DNA assembly |
| JP480 | actcaatataagcatatcccGCATAATCAAATGTTAGTTCAC | 5’ *P_1_-P_2_-*RCd12 Hifi DNA assembly |
| **pMTL84121 cloning** | | |
| JP481 | AGCAGGATCCTAACATATTTAGTATATACCTATGTATATATATTTAAAAC | 5’ *P-CD0977.1*-RCd11*-*  *P_2_-P_1_* BamHI |
| JP482 | AGCACTCGAGGAGCTTTTTAGGACAAAAAC | 3’ *P-CD0977.1*-RCd11*-*  *P_2_-P_1_* -XhoI |
| JP659 | AGCAGGATCCGCCGCCTCAATACTTATAAAG | 5’ *P-CD0956.2*-RCd10*-*  *P-* BamHI |
| JP660 | AGCACTCGAGCATTTTTTTTATATAAACAATGAAATTCAAG | 3’ *P-CD0956.2*-RCd10*-*  *P-* XhoI |
| JP661 | AGCAGGATCCATGTTTAAAATATATTGAATAGCTCTTG | 5’ *P-CD0904.1*-RCd13*-*  *P-* BamHI |
| JP662 | AGCACTCGAGATAAGACTGTGGAATATACAAAATTG | 3’ *P-CD0904.1*-RCd13*-*  *P-* XhoI |
| JP663 | AGCAGGATCCGTAGTTCTTCTTGCTTTTATTATACCAC | *5’-P-CD0956.3*-RCd14-*P*-BamHI |
| JP664 | AGCACTCGAGCAATTGTTAAACTTCAATTTATCG | *3’-PCD0956.3*-RCd14-*P*-XhoI |
| **pMSR and pMSR0 construction** | |  |
| JP416 | GGATCCTCTAGAGTCGACG | 5’ *codA* removal from pMTL-SC7215 and pMTL-SC7315 by inverse PCR |
| JP417 | GTAATCATGGTCATATGGATACAG | 3’ *codA* removal from pMTL-SC7215 and pMTL-SC7315 by inverse PCR |
| JP418 | atccatatgaccatgattacCGAATTCTGCATCAAGCTAG | 5' *P_tet_-CD2517.1* and 5’ *P_tet_-CD2517.1*-RCd8*-P* |
| JP419 | acgtcgactctagaggatccGAAGAACTACAATCTATTTTGC | 3' *P_tet_-CD2517.1* |
| JP619 | CCATACGATGTTCCAGATTAC | 3' *P_tet_-CD2517.1*-RCd8*-P* |
| **pMSR cloning** | |  |
| JP449 | CGTTTTGTAAACGAATTGC | 5’pMSR insert screening |
| JP450 | CTCACGTTAAGGGATTTTG | 3’pMSR insert screening |
| JP217 | ttttttgttaccctaagtttGAGAATTACTTTACAGATATAAAGAAAC | 5’ left arm Δ*CD0956.2* |
| JP218 | tgtagattacGGCTTCCACTTTAATTTTTGAATAC | 3’ left arm Δ*CD0956.2* |
| JP219 | agtggaagccGTAATCTACAATTTATAGAGTGGAG | 5’ right arm Δ*CD0956.2* |
| JP171 | gattatcaaaaaggagtttGCTCACTGCAAAATCTCTAC | 3’ right arm Δ*CD0956.2* |
| JP221 | GTATTCAAAAATTAAAGTGGAAGC | 5’ Δ*CD0956.2* screening |
| JP173 | CACACCCTCAATACCTTG | 3’ Δ*CD0956.2* screening |
| JP226 | ttttttgttaccctaagtttCCTAGTTTCTGCATACTTTC | 5’ left arm Δ*CD0977.1* |
| JP223 | gaatatattgGCTAGCTTTATTACATATATTATTTCCAG | 3’ left arm Δ*CD0977.1* |
| JP224 | taaagctagcCAATATATTCTATATTGCAATAAAAAACATTG | 5’ right arm Δ*CD0977.1* |
| JP156 | gattatcaaaaaggagtttGATAAGAACTATAAAGAGTTTATAAAATTATTAAAC | 3’ right arm Δ*CD0977.1* |
| JP163 | CATAATATCCCCTTCTTTCTATTAAC | 5’ Δ*CD0977.1* screening |
| JP164 | GCAGAAATAGAGGAAAGAGTG | 3’ Δ*CD0977.1* screening |
| JP523 | ttttttgttaccctaagtttGCAAGTGTTAGTGATATTACAG | 5’ left arm ΔψCD630_2 |
| JP524 | ataattatataaATGACTGCTTATAACAATATAGC | 3’ left arm ΔψCD630_2 |
| JP525 | aagcagtcatTTATATAATTATTTGGACTAACATATAGTATC | 5’ right arm ΔψCD630_2 |
| JP526 | agattatcaaaaaggagtttGGGGATTATCAGCAATGAAC | 3’ right arm ΔψCD630_2 |
| JP527 | GATAGTTATGGATTCTCATGGTG | 3’ ΔψCD630_2 screening |
| JP528 | CTTGTGTCTAACCTTTGCATC | 5’ ΔψCD630_2 screening |
| JP575 | gtgttttttgttaccctaagtttCAGGAGCTGACAATGTTC | 5’ left arm Δ*CD0904.1* |
| JP576 | ataaaaagttaatCCTATATATTACTTTTTTATTCATCCATATATC | 3’ left arm Δ*CD0904.1* |
| JP577 | aatatataggATTAACTTTTTATTGAGTATAGTAGC | 5’ right arm Δ*CD0904.1* |
| JP578 | agattatcaaaaaggagtttAATACCTTCAAATGAAAGAAGAG | 3’ right arm Δ*CD0904.1* |
| JP579 | GCCAGTCATTTCTTCTATGTACTC | 5’ Δ*CD0904.1* screening |
| JP580 | CGACATTGCTATGTCAAAAG | 3’ Δ*CD0904.1* screening |
| JP581 | gtgttttttgttaccctaagtttGGTTGATAATGTAAAATATGTAGC | 5’ left arm Δ*CD0956.3* |
| JP582 | ataaaaaaccCAATGAAATTCAAGTAAATAAATACC | 3’ left arm Δ*CD0956.3* |
| JP583 | aatttcattgGGTTTTTTATTAAGCATACTAGC | 5’ right arm Δ*CD0956.3* |
| JP584 | agattatcaaaaaggagtttCCAAGTGCTTTATCTATATTATTTATTC | 3’ right arm Δ*CD0956.3* |
| JP585 | CTTCACTCTACCGCAAAATAG | 5’ Δ*CD0956.3* screening |
| JP586 | GCGTTCTCAAACCTTTACATAC | 3’ Δ*CD0956.3* screening |
| JP596 | gtgttttttgttaccctaagtttGGCTATAGAGCAGCAAGAG | 5’ left arm, *P_thl_-ermB* insertion |
| JP597 | ctaaaccattCAATACAATTAATAAAGATTTGTACTACAC | 3’ left arm, *P_thl_-ermB* insertion |
| JP602 | ttgcttttatCTAATGTCAATGTAGGTATTCTTTG | 5’ right arm, *P_thl_-ermB* insertion |
| JP603 | agattatcaaaaaggagtttCCAGTTGTAGATATAGTTGG | 3’ right arm, *P_thl_-ermB* insertion |
| JP604 | GTCAAAAGATGAACATATAAGTAAATC | 5’ *P_thl_-ermB* insertion screening |
| JP606 | GCTATTGTATCTCTGTTTGAATCTAC | 3’ *P_thl_-ermB* insertion screening |
| JP673 | aaatctttattaattgtattgccaacgcgttatattgataaaaataataatag | 5’ *P_thl_-ermB*, *P_thl_-ermB* insertion |
| JP674 | aatacctacattgacattagcgtgcgactcatagaattatttc | 3’ *P_thl_-ermB*, *P_thl_-ermB* insertion |
| **PCR and quantitative PCR** | | |
| RT_polIII-F_Cdiff | TCCATCTATTGCAGGGTGGT | 5' CD1305 |
| RT_polIII-R_Cdiff | CCCAACTCTTCGCTAAGCAC | 3' CD1305 |
| JP353 | CTTTCGCCATCTTTTGATAGAA | 3’ flanking PhiCD630-1 |
| JP354 | TATGGTGTTGTGGCAATGAA | 5’ flanking PhiCD630-1 |
| JP352 | AAAATGCCGTGAAATCAATG | 5’ circular PhiCD630-1 |
| JP355 | CGCTAAAACTACACGTTTTAACATT | 3’ circular PhiCD630-1 |
| JP633 | AACCTCTCTTTGATGGCTCT | 5’ flanking PhiCD630-2 |
| JP634 | GGGGAATTTCTATGTTTACTCGG | 3’ flanking PhiCD630-2 |

Restriction sites are underlined.
