## Supplemental Data 3 for "Type I toxin-antitoxin systems contribute to mobile genetic elements maintenance in *Clostridioides difficile* and can be used as a counter-selectable marker for chromosomal manipulation"

**Table S3. The antisense RNA and associated toxin mRNA extremity identification for RCd11/RCd12-*CD0977.1/CD2889* by 5’/3’RACE.**

| **Name** | **Description** | **5’-end RACE position** | **5’-end TSS mapping position** | **Strand** | **3’-end RACE position** | **Size, nt** |
| --- | --- | --- | --- | --- | --- | --- |
| RCd11 (CD630_n00390) | Antitoxin of TA associated with cdi1_5 | 1142666,  1142673  1142439 | 1142666,  1142673  1142439 | - | 1142531,  1142295,  1142289 | 143, 151, 385 |
| CD0977.1 | Toxin of TA associated with cdi1_5 | 1142137 | 1142158 | + | 1142418 | 282 |
| RCd12 (CD630_n00980) | Antitoxin of TA associated with cdi1_4 | 3379972,  3377979,  3380206 | 3379972,  3377979,  3380206 | + | 3380346,  3380352,  3380114 | 143, 151, 385 |
| CD2889 | Toxin of TA associated with cdi1_4 | 3380505 | 3380505 | - | 3380223 | 282 |

The positions of 5’-start and 3’-end of these RNAs were identified by 5’/3’RACE analysis and compared with 5’ –end identified by 5’-end RNA-seq analysis (TSS mapping).
